# Genomic innovation through fragmentation and gene transfer underlies adaptation in pyrophilous bacteria

**DOI:** 10.1101/2025.11.11.687939

**Authors:** Ehsan Sari, Dylan J. Enright, Esbeiry S. Cordova-Ortiz, Maria E. Ordoñez, Andrew Byrd, Steven D. Allison, Peter M. Homyak, Michael J. Wilkins, Sydney I. Glassman

## Abstract

Wildfires reshape soil carbon (C) and nitrogen (N) dynamics by generating pyrogenic organic matter (PyOM) and selecting for pyrophilous “fire-loving” bacteria, yet the genomic mechanisms underpinning their post-fire adaptations are largely unknown. We combined comparative genomics of 16 pyrophilous bacteria spanning three phyla with bioassays and transcriptome profiling following PyOM exposure to test whether post-fire conditions promote genomic innovation within bacterial populations. Pyrophilous bacterial isolates exhibited genomic novelty through gene fragmentation, horizontal gene transfer (HGT), and plasmid-mediated operon expansion, enriching aromatic C degradation and N acquisition genes relative to non-pyrophilous sister taxa. Plasmid-encoded PyOM metabolism genes showed evidence of HGT across pyrophilous bacterial orders and cross-kingdom with pyrophilous fungi. Bioassays revealed overflow organic acid production associated with variation in tricarboxylic acid cycle genes, potentially moderating elevated post-fire soil pH and facilitating microbial succession. Our findings reveal how fire selects for bacterial genomic innovation and illuminate microbial evolution in extreme environments.

## Main

Wildfire frequency and severity are increasing globally^1^, profoundly altering microbiomes^2,3^ and the ecosystem functions they regulate^4^. Fires reduce microbial richness and biomass^5,6^, but promote the proliferation of pyrophilous “fire-loving” soil bacteria that were rare or absent pre-fire^7,8^, with implications for soil carbon (C) and nitrogen (N) cycling^9-11^. Fires also increase soil pH, combust and release inorganic N, and generate ash and pyrogenic organic matter (PyOM), a chemically complex and aromatic C-rich substrate^12-15^. Fires produce smoke, which may disperse and select for diverse pyrophilous bacteria adapted to survive in chemically complex aerosols^16^. Extreme abiotic stress caused by fire includes thermal shock, desiccation, and nutrient alteration^17,18^. Thus, post-fire environments combine strong selective pressure and novel resource availability to which pyrophilous bacteria must adapt.

Understanding the traits and genomics underpinning the evolution of pyrophilous bacteria is critical for elucidating their adaptive strategies, resilience, and contributions to post-fire ecosystem recovery, which have implications for global biogeochemical cycles. A mechanistic understanding is essential because how microbes survive and function post-fire is likely shaped by the strategies used to balance life history traits involving acquisition of post-fire resources, thermotolerance, and fast colonization^7,8^. Fire drives rapid microbial turnover and selects for stress tolerance and metabolic flexibility, making post-fire environments uniquely powerful systems for studying the genomic basis of microbial adaptation.

We^19^, and others^20,21^, have previously revealed mechanisms of genomic innovation and adaptation in pyrophilous fungi, including the trait trade-offs that enable rapid adaptation to burned soils, but genomic mechanisms are largely unknown in pyrophilous bacteria. Metagenomic surveys of burned soils suggest that pyrophilous bacteria are predominantly found within the phyla Actinomycetota, Pseudomonadota, and Bacillota, which harbor expanded aromatic C degradation and N acquisition pathways^22-24^. Yet, metagenome assembled genomes (MAGs) are typically insufficiently complete or contiguous to resolve the underlying genomic mechanisms^23^. Moreover, variation in the tricarboxylic acid (TCA) cycle pathway, which connects central metabolism to aromatic C turnover^25^, may also play a critical role in PyOM degradation. Thus, the evolutionary origins and functional dynamics of aromatic C degradation, N acquisition, and central C metabolism in pyrophilous bacteria remain unresolved.

Gene fragmentation, horizontal gene transfer (HGT), and plasmid-mediated operon expansion are key mechanisms by which bacteria generate genomic novelty^26^. Gene fragmentation can create modular or truncated genes that evolve new functions or regulatory properties^27^. HGT, which we previously showed spans ecological and phylogenetic boundaries within pyrophilous fungi^19^, facilitates genetic exchange within microbiomes, rapidly introducing new metabolic capabilities^28^. Plasmid-mediated operon expansion further enables the coordinated gain of multiple functionally linked genes, often conferring adaptive advantages under fluctuating or extreme environments^29^. Together, these processes underpin bacterial abilities to expand metabolism, exploit novel ecological niches, and respond to environmental pressures, including those imposed by fire. Identifying mechanisms of gene fragmentation, HGT, and plasmid-mediated operon expansion in pyrophilous bacterial genomes can reveal general principles of microbial adaptability and evolutionary innovation. More broadly, these mechanisms illuminate fundamental routes by which bacteria diversify and evolve across Earth’s varied environments.

Here, we test the overarching hypothesis that fire selects for genomic novelty in lineages of pyrophilous bacteria. We leveraged comparative genomics of 16 pyrophilous bacteria that we isolated from post-fire soils and smoke^30^, and conducted transcriptomics and bioassays on a subset of these isolates to investigate the evolution of aromatic C degradation, N acquisition, and TCA cycle genes. We show that genomic novelty in these genes arises through three distinct mechanisms—fragmentation, HGT, and plasmid-borne operon expansion—collectively enhancing genomic potential in pyrophilous compared to non-pyrophilous bacteria. We uncover mechanisms by which pyrophilous bacteria diversify their metabolism, alter soil chemistry, and support post-fire recovery, highlighting general principles and mechanisms of adaptive diversification among bacteria.

## Results

### Genome assembly

We sequenced genomes of 16 bacteria representing the three dominant pyrophilous phyla—Actinomycetota, Bacillota, and Pseudomonadota^8,18,31-33^—isolated from burned soils and smoke following Southern California wildfires^30^ (Table S1). We also downloaded the genomes of 48 sister isolates, selected based on their phylogenetic relatedness to the 16 pyrophilous isolates (Table S2). We generated ∼117 Gb of PacBio HiFi sequences equal to 37–358X genome coverage, achieving single contig assemblies ranging from 2.5 to 8.9 Mb (Table S1). Eight isolates had plasmids, with six isolates containing 1-3 plasmids, *Noviherbaspirillum soli* harboring six, and *Streptomyces longhuiensis* harboring four.

### Enrichment of aromatic C degradation and N acquisition genes in pyrophilous bacteria and underlying mechanisms

To test enrichment of genes in pyrophilous vs. non-pyrophilous bacteria, we focused on genes identified from previous post-fire soil metagenomic studies as critical for aromatic C degradation and N acquisition^18,23,24^ (Table S3). Compared to their non-pyrophilous sister taxa, we found significant enrichment (P < 0.05) of these genes in all but two pyrophilous bacterial isolates (*Bacillus pumilus* and *Modestobacter versicolor*) (Fig. 1). The pyrophilous isolates with the highest enrichment compared to their sister taxa that were not isolated from burned environments were *Pseudomonas* sp., which encoded 2X more orthologues for aromatic C degradation and N acquisition compared to its sister isolate *P. lutea* DSM 17257. Two isolates of *Peribacillus simplex*, P-BS and P-BF, encoded nearly double the aromatic C degradation and N acquisition orthologues as their sister isolate *P. simplex* WH6. The mechanisms underlying these gene enrichments in the pyrophilous isolates varied by taxa with presence/absence variation contributing most to *Pseudomonas* sp. and *N. soli*, and gene fragmentation contributing most to *P. simplex* P-BS and P-BF. Notably, gene fragmentation was ubiquitous across our pyrophilous isolates, with the only isolates not exhibiting gene fragmentation being the two without significant enrichment compared to their non-pyrophilous sister taxa (*B. pumilus* and *M. versicolor*), and *Streptomyces longhuiensis*, where presence-absence variation was a major driver of gene enrichment.

**Fig 1.**
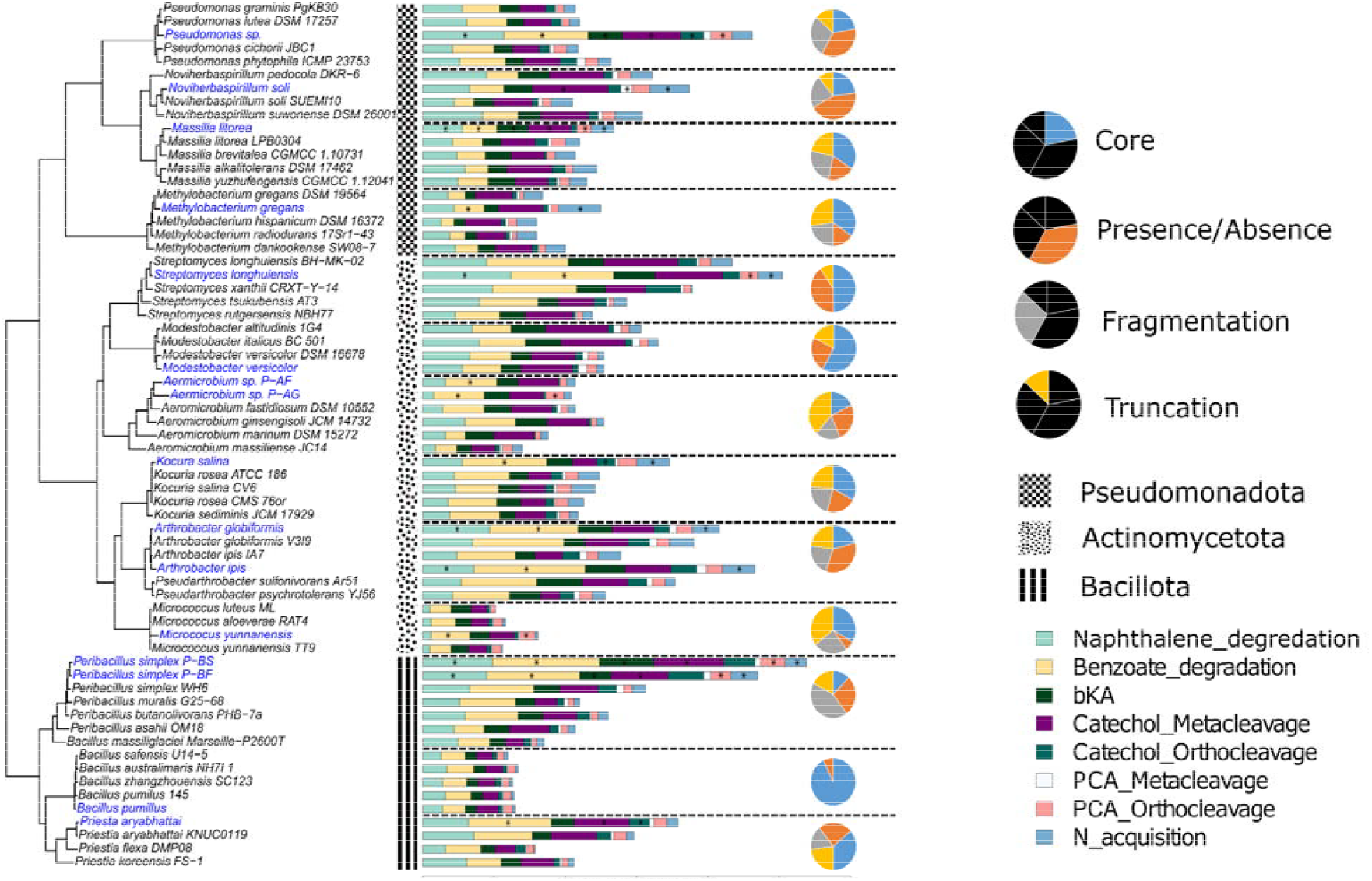
Strain-level enrichment of aromatic carbon (C) and nitrogen (N) acquisition genes and underlying mechanisms in pyrophilous bacteria. Orthofinder analysis of 16 pyrophilous bacteria that we isolated and sequenced (blue leaves on the tree) and 45 sister publicly available taxa (black) revealed patterns of gene enrichment in our isolates versus closely related taxa isolates from non-burned environments. The bar chart shows the total number of orthologues in each isolate with asterisks on bars showing significant difference (P < 0.05) between pyrophilous and non-pyrophilous sister isolates inferred from a Poisson distribution test, and the pie chart summarizes the mechanisms contributing to gene enrichment, inferred from comparisons between pyrophilous isolates and their sister taxa. Pyrophilous bacteria and their sister isolates are separated from loosely related species by dashed horizontal lines. Core genes were shared between pyrophilous bacteria and their sister isolates, presence/absence genes were unique to pyrophilous bacteria, fragmented genes appeared as multiple smaller gene segments, and truncated genes exhibited >20% reduction in amino acid sequence length relative to their orthologues. Abbreviations: bKA, beta-ketoadipate pathway; PCA, protocatechuate pathway; N_acquisition, nitrogen acquisition pathway.

### Transcriptome analysis after PyOM amendment

We conducted transcriptome analysis of two pyrophilous isolates with the highest enrichment of aromatic C and N acquisition genes, *Pseudomonas* sp. and *P. simplex* P-BS, before and at 1, 2, 4, and 16 h following PyOM exposure. In *P. simplex* P-BS, we identified 150 aromatic C degradation and N acquisition genes differentially transcribed (P< 0.05) after PyOM exposure; 49 showed differences across all time points, and 46 of these were upregulated (Supplementary File 1). In *Pseudomonas* sp., 90 aromatic C degradation genes were differentially transcribed (P< 0.05), with 51 remaining stable over time, and 28 upregulated. In both isolates, a gene encoding aminomuconate-semialdehyde (*dmpC_xylG_praB*) from the benzoate degradation pathway, exhibited the highest transcription change after PyOM exposure.

### Gene fragmentation drives genome novelty in pyrophilous *P. simplex* isolates

We focused on two *P. simplex* isolates, P-BS and P-BF, which exhibited the greatest putative gene fragmentation (Fig. 1), to examine how the reshuffling of the structural genes contributes to gene content differences relative to non-pyrophilous bacteria. We first compared genome-wide alignments between the two pyrophilous isolates (P-BS and P-BF) and their sister taxon WH6 isolated from unburned soil and found >0.99 average nucleotide identity (ANI) across the 3 genomes. P-BS had three major alignment gaps and P-BF had two when aligned to WH6 (Fig. 2A), which likely arose from large-scale rearrangements rather than nucleotide divergence. We cannot attribute these putative fragmentation events to overall sequence divergence given the high ANI between the pyrophilous isolates (P-BS and P-BF) and WH6 (Fig. 2A) and close phylogenetic distance (Fig. 1).

**Fig 2.**
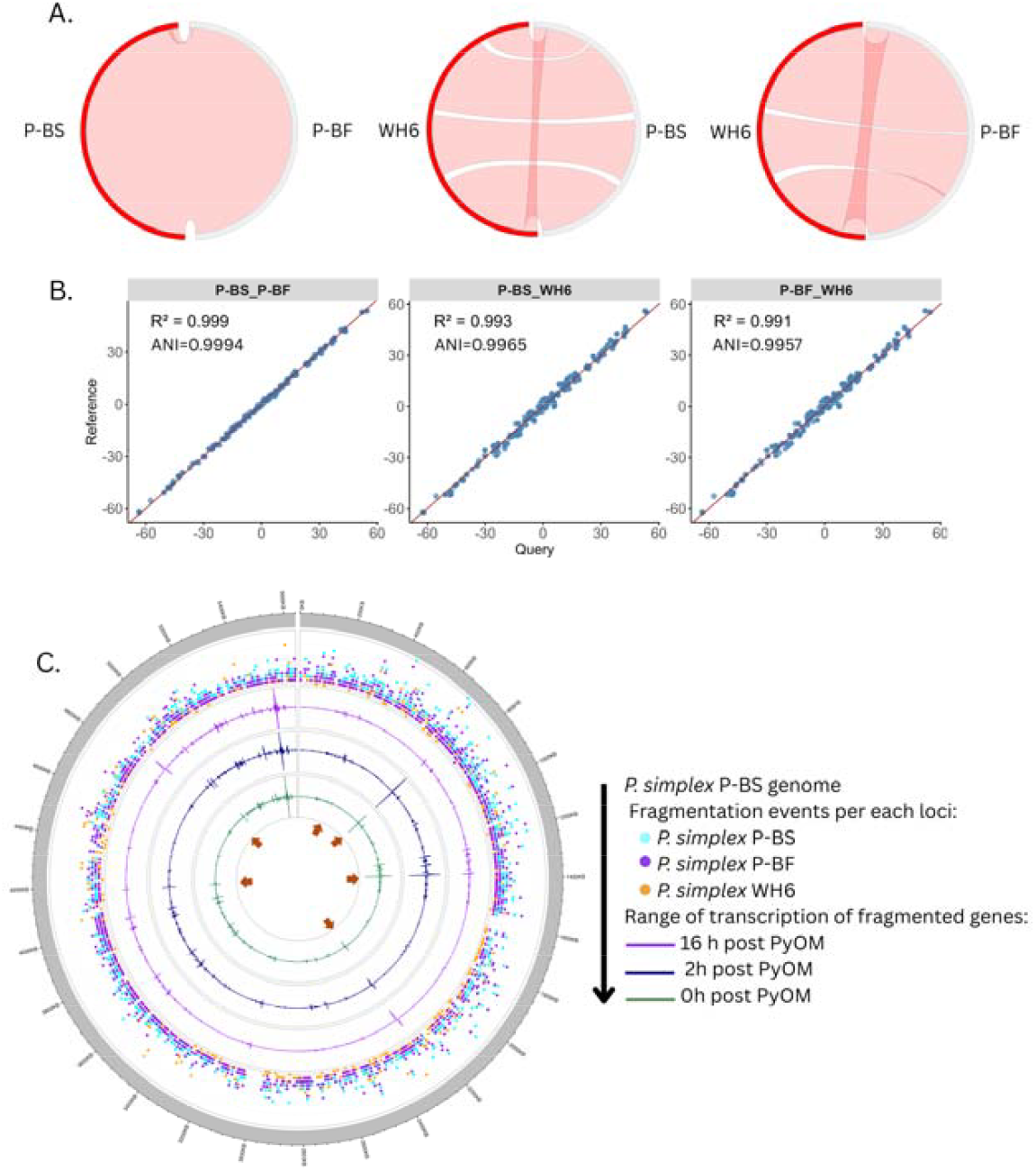
Genome-wide gene-fragmentation in pyrophilous *Peribacillus simplex* isolates (P-BS and P-BF) vs. non-pyrophilous isolate WH6 and the impact of fragmentation on the transcription of derivative genes. (A) Genome-scale alignment of P-BS and P-BF with WH6. Red and gray bars represent the genomes, and pink ribbons connect aligned regions. The white ribbons show three large-scale rearrangements in the P-BS-vs-WH6 alignment and two in the P-BF-vs-WH6 alignment. (B) Average nucleotide identity (ANI) and tetra-nucleotide frequency regressions between the genomes. Tetra-nucleotide regression measures similarity based on all possible 4-base combinations. R^2^ indicates the regression coefficient, and ANI values were calculated using JSpecies software. (C) Comparison of gene fragmentation events in P-BS and P-BF in reference to isolate WH6, highlighting the impact of fragmentation on the transcription of derivative genes. Brown arrows indicate examples of fragmented genes whose derivatives exhibit differential transcription after pyrogenic organic matter (PyOM) exposure. The gray circle represents the genome of isolate WH6, with ticks around the circle indicating positions along the genome in 20 -Kb intervals. The range of transcription of fragmented genes represent the genome-wide transcription per million (TPM) value of fragmented gene derivatives of isolate P-BS before (0h) or 2 and 16 h post PyOM exposure that show the highest transcription discrimination among samples.

We quantified the genome-wide distribution of putative gene fragmentation events in P-BS and P-BF compared to WH6 and paired these structural patterns with transcriptomic data after PyOM exposure to assess their regulatory consequences. Both P-BS and P-BF exhibited substantially higher fragmentation frequencies than WH6 across their genomes, consistent with a genome-wide process rather than isolated events (Fig. 2B). Some gene-fragment derivatives were differentially transcribed following PyOM exposure in P-BS (Fig. 2B, brown arrows), indicating that some fragmentation events have regulatory or functional relevance. Unlike *Pseudomonas* sp., where presence/absence variation is the primary source of genome novelty (Fig. 1), fragmented derivatives in P-BS showed greater transcriptional variation after PyOM exposure in selected aromatic C degradation and N acquisition genes (Table S2; Fig. S1A). To explore mechanisms underlying this variation, we performed motif-enrichment analysis on intergenic regions separating fragmented ORFs in P-BS. The five most enriched motifs matched four transcription factor binding sites and one RNA polymerase σ70 site (Fig. S1B), suggesting that regulatory elements introduced at fragmentation boundaries contribute to the differential transcription of these gene segments.

The aromatic C degradation gene encoding *dmpC_xylG_praB*, which showed the highest PyOM-responsive transcription in both P-BS and *Pseudomonas*, also displayed clear structural disruption evidenced by reduced proteome sizes across 11 of its 22 orthogroups (Fig. S2A). We focused on these orthogroups in P-BS and P-BF to understand how fragmentation shapes gene-content differences relative to the non-pyrophilous isolates. A unique gene duplication event within the HOG0000303 orthogroup occurred in *P. simplex* WH6, with one copy representing the ancestral full-length version and the corresponding copies in P-BS and P-BF exhibiting fragmentation (Fig. S2B). Of the four resulting fragmented derivatives in P-BS, two of them were differentially transcribed following PyOM exposure (Fig. S2C), providing further support that fragmentation creates heterogeneity in regulatory behavior among gene segments.

### HGT drives genomic novelty in the pyrophilous bacterium *N. soli*

Gene presence/absence variation was the dominant driver of gene-content differences between *N. soli* and its non-pyrophilous sister isolates (Fig. 1). The presence of six candidate plasmids (5–785 kb; Table S1) in *N. soli* suggested a role for plasmid-mediated HGT in shaping gene variation. Sequencing read coverage for the three largest plasmids was comparable to the median genome coverage (Fig. S3), indicating stable maintenance and likely functional relevance in *N. soli* metabolism. Three of the six plasmids carried Type IV secretion system genes, suggesting conjugative capacity and direct plasmid-mediated HGT (Fig. 3A).

**Fig 3.**
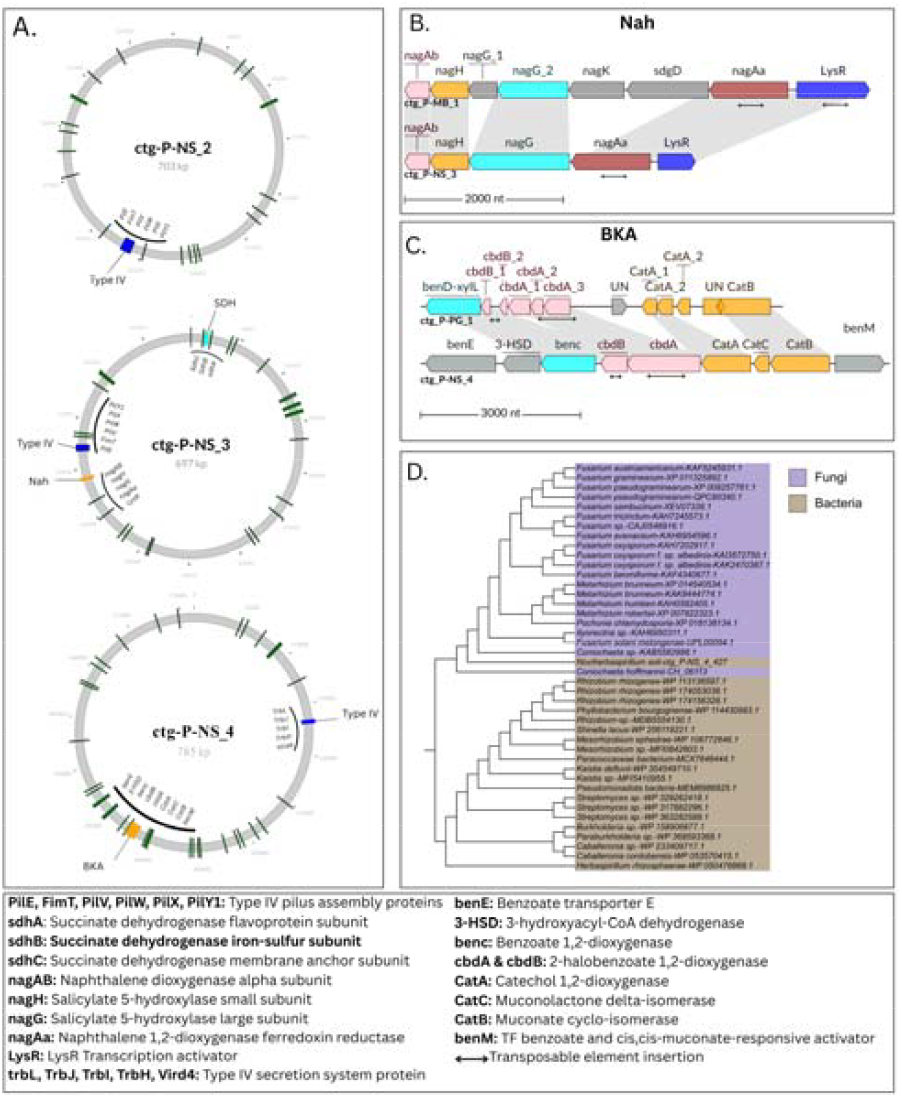
Conjugative plasmids in *Noviherbaspirillum soli*, showing evidence of horizontal gene transfer (HGT) into pyrophilous bacteria and cross-kingdom transfer into pyrophilous fungi. (A) Circular map of the three largest conjugative plasmids in *N. soli*. Circles represent three plasmids: *ctg_P-NS_2, ctg_P-NS_3*, and *ctg_P-NS_4*. Green ticks on circles indicate genes encoding tricarboxylic transport membrane proteins, blue ticks mark Type IV secretion system genes, orange ticks indicate aromatic C degradation genes involved in naphthalene degradation (Nah) and beta-ketoadipate (BKA) pathways, and cyan tick indicates a succinate dehydrogenase operon (SDH) encoding TCA cycle genes. The Type IV genes on *ctg_P-NS_2* and *ctg_P-NS_3* were identical, including pili-making genes *PilE, FimT, PilV, PilW, PilX, PilY2*, while those on *ctg_P-NS_4* belonged to the *tra/trb* operon including the pili-making genes *TrbL, TrbJ, TrbI, TrbH* and the DNA/protein secretion gene, *VirD4*. Putative HGT into pyrophilous bacteria *Massilia litorea* (B) and *Pseudomonas* sp. (C) and cross-kingdom transfer into pyrophilous fungus *Coniochaeta hoffmannii* (D). Pentagons in B & C represent plasmid genes (bottom) and their corresponding genomic introgression sites. Colors of the genes and connecting ribbons indicate sequence homology between plasmid and genomic regions. In *M. litorea*, the *nagG* gene appeared fragmented and diverged; among the two tandem *nagG* copies (*nagG-1* and *nagG-2*), only *nagG-1* retained homology with the plasmid gene (B). In *Pseudomonas* sp., BKA genes (*cdbA, cdbB, catA*) showed fragmentation co-localized with transposable element insertions (C). Panel C shows phylogeny of catechol-1,2-dioxygenase (*CatA*) in *N. soli* (ctg_P-NS_4_427) vs. its orthologues in bacteria (brown) and fungi (purple) identified via NCBI BLASTP. The tree also includes a gene (CH_06113.t1) encoding *CatA* horizontally transferred into pyrophilous fungus *Coniochaeta hoffmannii*^*19*^. Tree leaves are labeled with the species name followed by the UniProt ID of the BLASTP hit.

Two of these plasmids carried aromatic C degradation gene clusters, including a naphthalene degradation (Nah) and a β-ketoadipate (BKA) pathway operon. Notably, we detected putative transfer of re-arranged Nah and BKA operons from *N. soli* (Order Burkholderiales) plasmids into the genomes of *Massilia litorea* (Order Burkholderiales; Fig. 3B) and *Pseudomonas* sp. (Order Pseudomonadales; Fig. 3C). Transcription analysis revealed that only one of 12 BKA genes transferred into the *Pseudomonas* sp. genome was differentially transcribed after PyOM exposure (Fig. S4), suggesting that not all transferred genes are immediately co-opted into PyOM-responsive regulation.

A phylogenetic analysis of the plasmid’s catechol-1,2-dioxygenase (*catA*) genes in the BKA operon in concert with our previously analyzed genomes of pyrophilous fungi^19^, confirmed closest homology between the *N. soli* plasmid and a candidate HGT gene in pyrophilous fungus *Coniochaeta hoffmanii* (Fig. 3D). While this relationship does not resolve transfer directionality, the shared sequence similarity and phylogenetic coherence support the involvement of cross-domain HGT in the evolution of PyOM degradation pathways.

Additional resource acquisition genes on these plasmids included a succinate dehydrogenase (SDH) operon, encoding a core TCA cycle enzyme complex (Fig. 3A). We also identified multiple copies (22, 27, 37 per plasmid) of tricarboxylic transport membrane genes across the three plasmids of *N. soli*. These features suggest that plasmids contribute to variation in central C metabolism through TCA cycle in *N. soli*, although the functional consequences require further investigation.

### Plasmid-mediated operon expansion drives genomic novelty in *S. longhuiensis*

*Streptomyces longhuiensis* exhibited the second highest presence/absence variation among pyrophilous bacterial isolates examined (Fig. 1) and had the second most plasmids (4 plasmids size 7 to 24 kb; Table S1, Fig. 4A). None of the plasmids showed sequence homology with the genome of the non-pyrophilous sister isolate *S. longhuiensis* BH-MK-02 (Fig. S5), suggesting they are unique to the pyrophilous isolate.

**Fig 4.**
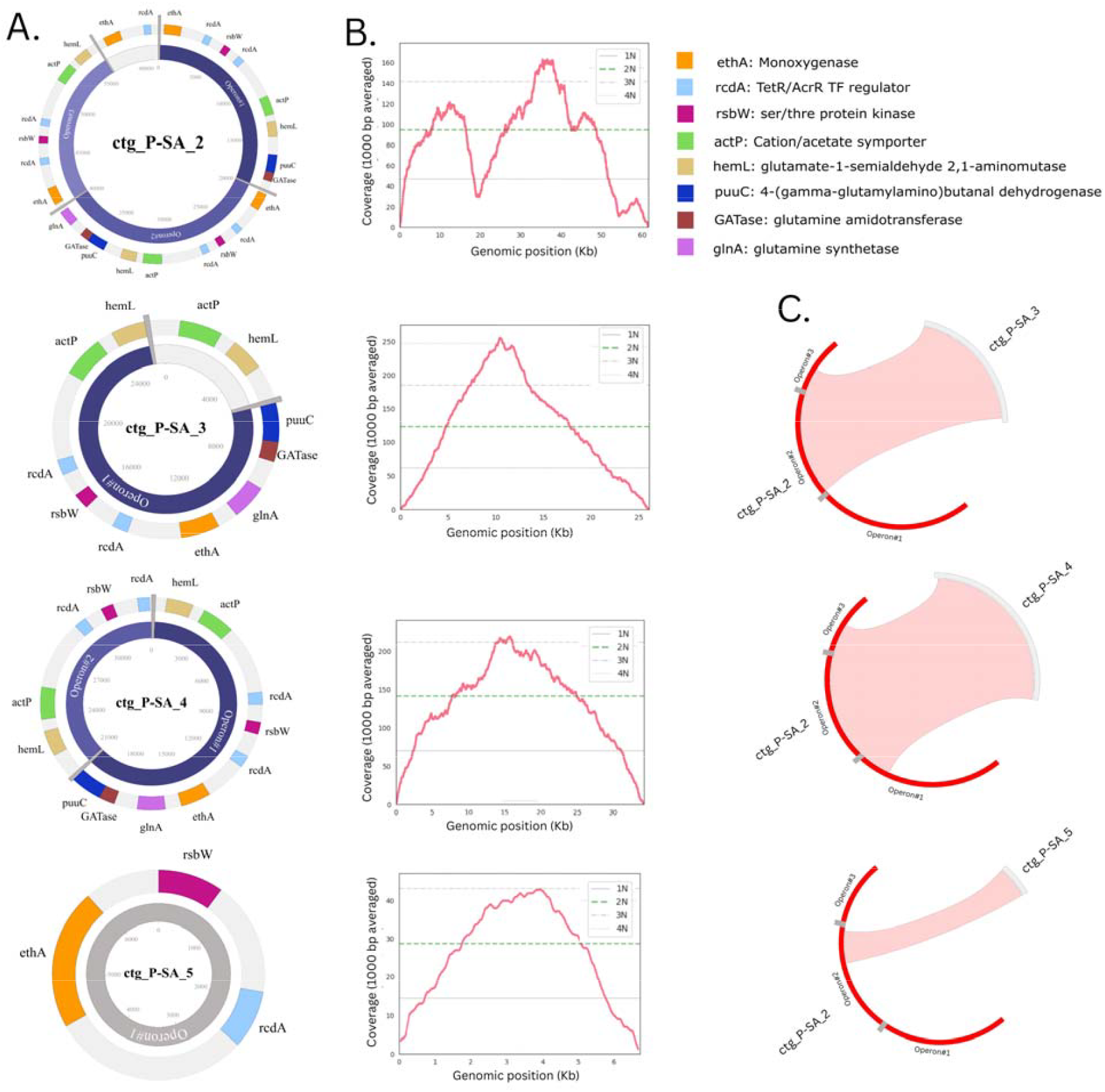
Glutamine synthesis operon on the plasmids of *Streptomyces longhuiensis*. (A) Plasmid maps harboring the operons. Multiple occurrences of the same operon are labeled as operon# on each plasmid. (B) Sequencing read coverage for each plasmid. Horizontal lines indicate coverage levels divided into four quartile bins (1-4N), and the green dashed line marks the median coverage. (C) Sequence alignment of *ctg_P-SA_2* against *ctg_P-SA_3-5* to evaluate whether *ctg_P-SA_2* could have arisen from a combination of other plasmids. Curved bars represent plasmid DNA, and pink ribbons connect aligned regions.

Each plasmid carried a glutamine synthetase operon, present in its most complete form as a nine-gene cluster, with 1 to 3 copies per plasmid (Fig. 4A). Sequence coverage was higher for the three plasmids than for the chromosomal contig, indicating the presence of multiple plasmid copies per cell (Fig. 4B). Coverage plots for three plasmids showed single peaks, marking putative origins of replication, whereas one plasmid exhibited at least two distinct peaks. Although this pattern might indicate plasmid fusion, sequence alignments showed no evidence that this plasmid resulted from recombination between the existing plasmids in the cell (Fig. 4C). Hence, the fusion-like event may have occurred prior to plasmid acquisition by *S. longhuiensis*.

Because all 4 plasmids lacked identifiable conjugation machinery (Fig. 4A), we examined whether plasmid-derived operon sequences had been incorporated into the *S. longhuiensis* chromosome. Protein-level comparisons of the operon structural genes, glutamine synthetase (*glnA*), revealed two chromosomal copies clustered closely with the plasmid-borne sequences (Fig. S6A). Further inspection identified a region consistent with the integration of a nearly intact plasmid-derived operon (Fig. S6B), adjacent to a well-characterized bacterial insertion sequence encoding a IS256 transposase, suggesting a model in which transposon-associated integration mediated operon expansion within the chromosome.

### TCA cycle variation drives acid production by pyrophilous bacteria

Several lines of evidence prompted us to examine the role of TCA cycle variation in pyrophilous bacterial adaptation to the post-fire environment. The TCA cycle oxidizes aromatic C intermediates to generate energy and biosynthetic precursors^25,34^. Conjugative plasmids of *N. soli* encoded multiple copies of tricarboxylic acid transporters and an operon containing *SDH*, a core TCA cycle gene (Fig. 3A). *Streptomyces longhuiensis* carried glutamine synthetase operons on plasmids that channel β-keto acid intermediates from the TCA cycle toward N assimilation^35^ (Fig. 4A). Finally, we observed differential transcription of 9 TCA cycle gene families in *P. simplex* P-BS (Fig. S7A) and 12 in *Pseudomonas* sp. (Fig. S7B) after PyOM exposure.

*P. simplex* P-BS and P-BF lacked functional copies of isocitrate dehydrogenase (*IDH*), which was otherwise present in all other isolates, including their non-pyrophilous sister isolate WH6 (Fig. 5A). Further analysis suggested that both P-BS and P-BF retained a significantly shorter homolog of *IDH*. Alignment with *IDH* sequences from 18 publicly available *P. simplex* genomes confirmed this smaller protein size being unique to P-BS and P-BF. Indeed, public genomes had ∼420 amino acids, whereas P-BS had 188 and P-BF had 109 amino acids (Fig. 5B).

**Fig 5.**
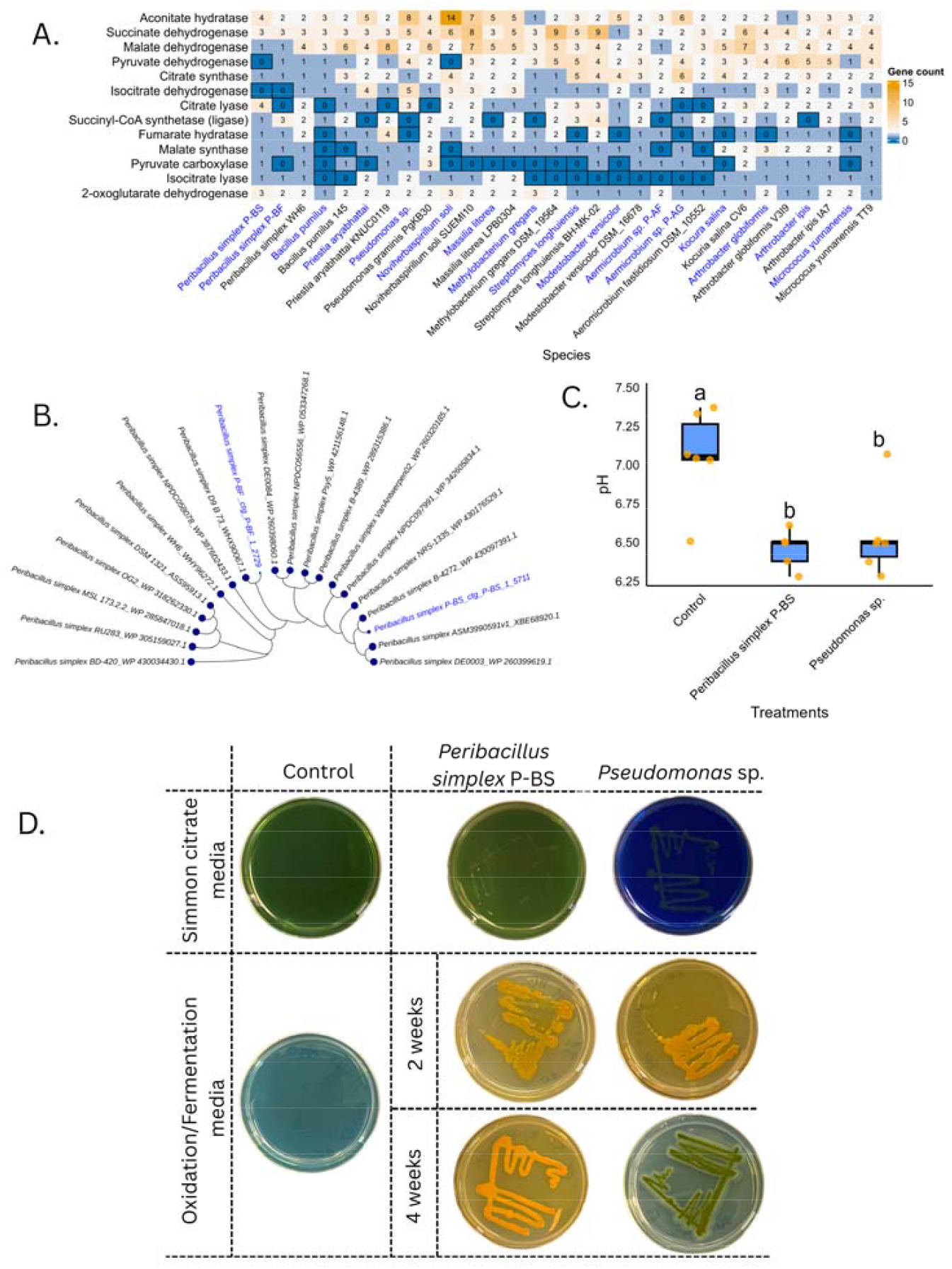
Genetic variation in the tricarboxylic acid (TCA) pathway between pyrophilous and non-pyrophilous isolates, highlighting a fragmented non-functional copy of isocitrate dehydrogenase (*IDH*) gene in the pyrophilous *Peribacillus simplex* P-BS associated with organic acid accumulation and potential acidification of the surrounding environment. (A) Number of gene copies in TCA pathway in pyrophilous bacteria and their closest sister isolates. Absence of genes is indicated by dark blue bordered cells. Taxa names in blue denote pyrophilous bacteria sequenced in this study. (B) Phylogeny of *IDH* proteins in publicly available *Peribacillus simplex* genomes compared with pyrophilous isolates P-BS and P-BF with fragmented non-functional copy of *IDH*. Blue tree leaves correspond to putative *IDH* proteins in P-BS and P-BF, and circles at branch tips represent amino acid sequence length. (C) Acidification of oxidative-fermentative (OF) media by P-BS and *Pseudomonas* sp. (acidifying pyrophilous isolate with functional *IDH*). The box plot shows pH in OF liquid media for P-BS and *Pseudomonas* sp., and the uncultured control; letters indicate groups with significantly different means (Tukey’s test, P < 0.05). (D) pH changes in Simmon’s citrate and oxidative/fermentative (OF) media by *Peribacillus simplex* P-BS and *Pseudomonas* sp. pH changes are shown relative to control plates with no bacteria cultured on them. In Simmon’s citrate media, the transition from green to blue indicates pH reduction due to citrate metabolism. In OF media, the transition from blue to yellow reflects media acidification, while a shift back from yellow to blue indicates pH increase as acids are later metabolized. The transition of Simmons’ citrate medium inoculated with *Pseudomonas* sp. indicated active citrate utilization, whereas no such transition was observed for P-BS. Additional evidence for citrate consumption by *Pseudomonas* sp., but not by P-BS, came from four-week-old OF medium cultures, in which only *Pseudomonas* sp. reverted the medium’s blue coloration.

Further analysis revealed that the *IDH* underwent a fragmentation event in P-BS, with three adjacent fragments corresponding to distinct segments of the full length *IDH* in WH6 (Fig. S8). We identified a 25-nucleotide insertion at the fragmentation point between the first two fragments, and a single-nucleotide polymorphism introducing a start codon between second and third gene fragments. These results indicate that isolates P-BS and P-BF experienced gene fragmentation events that likely generated non-functional copies of *IDH*, which might indicate an interrupted TCA cycle.

### Bioassay to test the impact of variation in TCA cycle on organic acid production and consumption

We used oxidation-fermentation (OF) media to assess whether IDH fragmentation in P-BS altered TCA mediated organic acid production. After 48h, both the *IDH-*fragmented P-BS and the *IDH-*intact *Pseudomonas* sp. significantly (P<0.05) reduced the media pH compared to the control (Fig. 5C), suggesting that multiple metabolic routes can generate overflow organic acids in pyrophilous bacteria. However, *Pseudomonas* sp. but not P-BS consumed organic acid (citrate in Simmon’s citrate media) as a C source (Fig. 5D), suggesting that in *Pseudomonas* sp., organic acids may also serve as an energy source. Further support came from four-week-old OF media, in which *Pseudomonas* sp. restored pH to neutral, suggesting the acid was later consumed, likely after glucose was depleted. In contrast, P-BS OF plates remained acidic, suggesting an inability to obtain energy from citrate.

## Discussion

This study presents the first whole-genome comparative genomic analysis of pyrophilous bacteria, previously known only from incomplete or fragmented MAGs^18,23^. Leveraging our large and diverse culture collection of pyrophilous bacteria (Table S1)^30^ and integrating genomics, transcriptomics, and targeted bioassays, we develop a conceptual model showing how gene fragmentation, plasmid-mediated operon expansion, and cross-order and cross-domain HGT lead to adaptive metabolic traits that promote survival and diversification in pyrophilous bacteria (Fig. 6). These genomic innovations reshape aromatic C degradation, N acquisition, and central C metabolism, including the TCA cycle, conferring metabolic flexibility under fire-altered nutrient and chemical conditions and positioning post-fire systems as powerful natural models for studying bacterial evolution and adaptive diversification.

**Fig 6.**
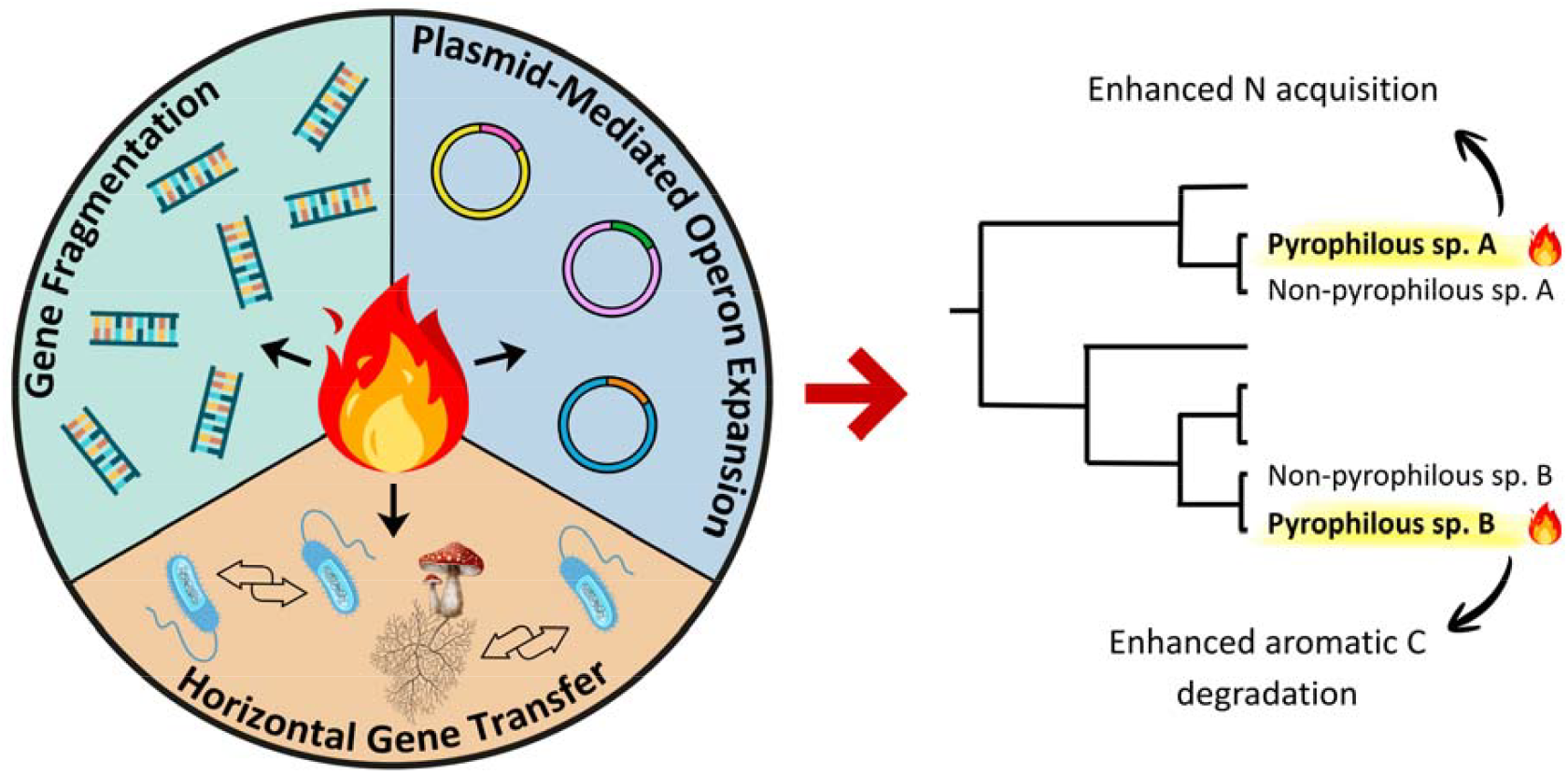
Conceptual model illustrating how fire selects for genomic novelty strategies that generate genomic diversity within bacterial lineages and enhance aromatic carbon (C) and nitrogen (N) acquisition gene repertoires in pyrophilous (isolated from burned soil or smoke) versus non-pyrophilous strains (isolated from non-burned environments) of the same species.

We found that gene fragmentation is a major contributor to genomic novelty, increasing enrichment of aromatic C degradation and N acquisition genes in 13 out of 16 of our pyrophilous bacterial isolates. Although gene disruption can reflect pseudogenization^27^, here we found that fragmented genes were transcribed and thus likely repurposed for regulatory behavior rather than becoming non-functional. The highest gene fragmentation occurred in *P. simplex* isolates P-BS and P-BF (Fig. 1), where differential transcription, motif enrichment, and repeated pattern across isolates (Figs. 2, S1, S2) suggest that these fragmented gene derivates are actively maintained. Fragmented loci may act as evolutionary intermediates by promoting exon shuffling, gene fusion/fission events, and recruitment of accessory genes, allowing pyrophilous microbes to fine-tune metabolic pathways under fire-induced selection pressures^27,36^. In stress-adapted microbial lineages, genome streamlining has been previously proposed to facilitate rapid genomic plasticity and regulatory rewiring in response to environmental stressors (e.g., nutrient limitation, oxidative stress, or thermal disturbance)^37-39^, suggesting that gene fragmentation may represent an adaptive response to stressful post-fire environments. Moreover, genes derived from fragmentation harbored transcription factor–binding sites and intergenic regulatory elements, likely explaining the variable transcription of fragmented derivatives after PyOM exposure (Figs. S1A, S2C). Overall, these findings highlight gene fragmentation as a key mechanism shaping genomic innovation and adaptive capacity in pyrophilous bacteria.

Plasmids were key contributors to genomic novelty in pyrophilous bacteria, carrying genes that expand post-fire aromatic C degradation and N acquisition. We found that conjugative plasmids containing nah and BKA pathway genes moved from *N. soli* to other Pseudomonadota genera, both within Burkholderales (*Noviherbaspirillum, Massilia*) and across orders to the Pseudomonadales (*Pseudomonas*) (Fig. 3). This finding is consistent with previous studies showing that conjugative plasmids have encoded aromatic C degradation functions and facilitated their transfer among soil bacteria^40-42^. Although notable within anaerobic gut fungi^43^, cross-domain HGT is not frequent. Here, we found that the *N. soli* plasmid gene catechol-1,2-dioxygenase (*catA*) clustered closest to the candidate *catA* in the fungus *Coniochaeta hoffmannii*, which we previously sequenced^19^, suggesting that *catA* might be horizontally transferred across domains of life. Interestingly, both the isolates of *N. soli* and *C. hoffmannii* that we characterized were isolated from the same soil sample at the same timepoint from the El Dorado fire burn scar, lending support to our findings suggesting cross-domain HGT.

Plasmid-mediated operon expansion also appeared to be a distinct mechanism of genomic innovation in some pyrophilous bacteria, as evidenced by the integration of the glutamine synthetase operon into the genome of *S. longhuiensis*. The structural gene within this operon, *glnA*, encodes a glutamine synthetase which is central to bacterial N assimilation and C-N metabolic integration by incorporating ammonium into glutamine and linking N assimilation to TCA cycle intermediates^35,42,44^. The observation that the plasmids had threefold higher read coverage than the genome indicates that *S. longhuiensis* maintains high-copy plasmids for advantages such as increased gene dosage (an increase in the number of copies of a particular gene, leading to elevated expression), enhanced plasmid stability, and a greater capacity for rapid adaptation^35,44^. Together, these results show that plasmids are central drivers of metabolic innovation in pyrophilous bacteria, enhancing evolutionary flexibility and expanding their capacity for aromatic C degradation and N acquisition in fire-impacted soils.

Variation among pyrophilous bacteria in TCA cycle genes, which play key roles in central C metabolism^25,34^, and are important intermediaries in PyOM breakdown pathways, may influence post-fire soil pH by driving organic acid production. Intra- and inter-species variation in TCA cycle gene complements is well documented in bacteria^34,45^, underscoring the metabolic plasticity of central C metabolism. These variations can lead to incomplete oxidation of TCA intermediates, resulting in the secretion of organic acids such as citrate, succinate, and fumarate^46^, aligning with our observation for *P. simplex* P-BS and *Pseudomonas* sp. Here, we found variation in TCA cycle genes, including a unique nonfunctional copy of *IDH* arising from two fragmentation events, was associated with organic acid overproduction and media acidification in pyrophilous bacteria *P. simplex* P-BS (Fig. 5). While we cannot prove causality, our phenotypic assays suggest that gene fragmentation may cause metabolic rerouting and organic acid secretion, which may allow pyrophilous bacteria to modify pH, which is often increased post-fire^12,15^. Our findings thus suggest that variation in central C metabolism of pyrophilous bacteria could influence soil chemistry following fire, potentially altering post-fire microbial succession and ecosystem recovery.

## Conclusion

We present the first comprehensive genomic analysis of pyrophilous bacteria, uncovering three key mechanisms—gene fragmentation, HGT across bacterial orders and cross-domain to pyrophilous fungi, and plasmid-mediated operon expansion—that drive adaptive metabolic traits to promote survival and diversification in pyrophilous bacteria (Fig. 6). These results underscore the role of genomic innovation in shaping microbial survival and function under fire-altered conditions, and build from previous literature showing that stressful conditions drive bacterial evolution via gene fragmentation^38,39^. Furthermore, we show that variation in central C metabolism gives rise to distinct pyrophilous traits, such as organic acid secretion that modulates post-fire soil chemistry, potentially lowering pH, with implications for global biogeochemical cycling. Collectively, our findings provide a framework for understanding how microbes evolve to persist and diversify in fire-prone and rapidly fluctuating environments.

## Materials and Methods

### Bacterial isolates

We leveraged our collection of pyrophilous bacteria isolated from post-fire soils and smoke across California^30^ and selected a subset of 16 pyrophilous isolates from 13 genera and three phyla (Table S1). We selected these isolates because they were abundant across a variety of fire-prone ecosystems^18,31-33^ and we expected them to exhibit a diversity of pyrophilous trait strategies^8^. We isolated 12 strains from soil slurries as published^30^. We isolated four strains from smoke by attaching petri dishes filled with Lysogeny Broth (LB) (Fisher Scientific, CAS# 154 73049-73-7) media to meter sticks, and holding them open for 20 minutes to collect ash from smoke during the peak of fire (fire names and locations in Table S1). All isolates were stored in glycerol stocks in the -80 °C and then Sanger sequenced to check identification upon reviving to ensure they were pure cultures.

### High Molecular Weight (HMW) DNA extraction and PacBio HiFi sequencing

We grew purified bacterial isolates in LB broth at 23 °C for 24-72 h based on their growth rate prior to extracting HMW DNA with Qiagen Genomic Tip 100/G (Qiagen, Hilden, Germany) following the manufacturer’s protocol. We evaluated DNA purity and concentration with the Qubit™ Fluorometer dsDNA BR Assay Kit (Invitrogen, Carlsbad, USA) using a DeNovix spectrophotometer Model Ds-11 (DeNovix Inc. Wilmington, USA). We measured DNA integrity and fragment size with the Genomic DNA Screen Tape (Agilent, Waldbronn, Germany) at the University of California, Riverside Institute for Integrative Genome Biology (UCR IIGB).

We then submitted DNA to the University of California Irvine Genomics Research and Technology Hub (UCI GRTH) for HiFi SMRT Bell Library Preparation and PacBio HiFi Sequencing. The sequencing library was prepared using SMRTbell template prep kit version 3.0 with overhang barcoded adapters (PacBio, Menlo Park, USA) and sequenced on the Sequel IIe™ System (PacBio, Menlo Park, USA). Samples were multiplexed to achieve the genome coverage ≥ 30X.

### RNA extraction and sequencing

We chose two pyrophilous bacteria, *P. simplex* P-BS and *Pseudomonas* sp., with the highest enrichment of aromatic C and N acquisition genes and ability to acidify media for transcriptome analysis. The isolates were grown in LB media for 24 h and the cells were pelleted by centrifuging at 3000 rpm for 3 minutes then washed twice with sterile distilled water to remove excess LB media. We then added custom-prepared PyOM media prepared from wildfire charcoal^30^ and harvested cells right before adding PyOM (0h), then 1, 2, 4, and 16 h after PyOM exposure. We extracted total RNA from the 30 samples (2 isolates × 3 replicates × 5 timepoints). For *P. simplex* P-BS, we used the E.Z.N.A.® bacterial RNA Kit (Omega Bio-Tek, Norcross, U.S.A.) following the manufacturer’s protocol. For *Pseudomonas* sp., we extracted total RNA using TRIzol^47^ as the kit did not yield enough RNA for sequencing. We then submitted RNA samples to the Department of Energy Joint Genome Institute in Berkeley, California for library preparation and sequencing, where mRNA libraries were prepared using the Illumina TruSeq® RNA Sample Preparation v. 2 Kit (Illumina, San Diego, USA) and sequenced on the NovaSeq 6000 system (Illumina, San Diego, USA).

### Genome assembly and annotation and transcription analysis

We generated genome assemblies for each isolate using Hifiasm v.0.16.1-r375^48^ with -l 0 parameter to disable purging haplotigs. We obtained the assembly metrics using QUAST software v.5.2.0^49^, and used BUSCO software v.5.4.3 to assess assembly completeness^50^. We annotated bacterial genomes using the Distilled and Refined Annotation of Metabolism (DRAM) pipeline^51^. Protein-coding sequences were functionally annotated using eggNOG-mapper to assign ortholog-based annotations and predict metabolic functions^52^.

We trimmed the sequencing adapters from RNA-seq Illumina reads with TrimGalore v.0.6.10 (https://github.com/FelixKrueger/TrimGalore), and filtered to retain only reads with a Phred quality score > 30 and length of ≥ 50 bp. We aligned the filtered reads to their corresponding genomes using STAR^53^ with --alignIntronMax 1 to suppress splicing. Normalized gene counts at 1, 2, 4, and 16 h after PyOM exposure were compared to the 0 h baseline using DESeq2^54^ and Log_2_ fold changes were calculated. Genes with adjusted P ≤ 0.05 were considered differentially transcribed. We excluded the 1-hour samples for *Pseudomonas* from data visualization due to high variation among replicates.

We generated 60.7 M 150-bp paired-end Illumina RNA-seq reads for *Pseudomonas* sp. and 51 M for *P. simplex* P-BS. After mapping, 93–96% of reads aligned uniquely to the *Pseudomonas* sp. and 92–95% to the *P. simplex* P-BS genome.

### Annotation of Aromatic C and N-acquisition and TCA cycle Genes

We used Hidden Markov Model (HMM) profiles for 27 aromatic C, 4 N acquisition, and 13 TCA cycle genes (Table S1) generated previously^55,56^. Aromatic C degradation genes examined belonged to meta- and ortho-cleavage for both catechol and protocatechuate and naphthalene and benzoate degradation pathways^22,56,57^. According to previous studies, bacteria transform aromatic C into intermediates such as catechol, protocatechuate, and benzoyl-CoA, followed by conversion to secondary intermediates such as acetyl-CoA, succinyl-CoA, and pyruvate through the β-ketoadipate (bKA) pathway^56,58^, so we also examined genes from the BKA pathway. We examined N acquisition genes such as nitrate and nitrite reductase whose transcription was previously shown to increase after fire^59^. We also included nitrous oxide reductase (*nosZ*) since high-severity fires may impair the soil microbiome’s capacity to convert nitrous oxide (N_2_O), a potent GHG, to nitrogen (N_2_) gas^60^, leading to elevated N_2_O emission from burnt soil^10^. We also examined TCA cycle genes due to their key role in central C metabolism^25,34^ and evidence found in this study for their contribution to the adaption of pyrophilous bacteria to post-fire environmental factors such as acidification.

### Orthogroups construction and visualization

We performed comparative genomic analysis of the 16 pyrophilous bacteria (Table S1) with 48 sister isolates (Table S2). To obtain the sister isolates, we downloaded the protein FASTA files of all publicly available genomes of each bacterial species using the get_assembly tool v.0.10.0 with the species name specified in the “organism” option (https://github.com/davised/get_assemblies). We then constructed a phylogenomic tree for each individual species using alignment generated with the BUSCO_phylogenomics tool release 20240919 (https://github.com/jamiemcg/BUSCO_phylogenomics). We constructed the phylogenomic tree from the trimmed alignment using FastTree v2.1.11^61^ software with -spr 4, -mlacc 2, -slownni, and -gamma parameters. We assigned the sister isolates to be those closely clustered with the target genomes. We assigned genes to orthogroups (clusters of orthologues across genomes) using Orthofinder v.2.5.5^62^ and used the gene IDs of HMMER annotated aromatic C and N acquisition and TCA cycle genes to extract assigned orthogroups from the Phylogenetic Hierarchical Orthogroups predicted for the oldest common ancestor of all species (N0). We obtained the orthogroups per gene family and individual genes by creating a matrix of orthologue counts per species. We visualized the sum of genes for each of naphthalene and benzoate degradation, BKA, catechol meta- and ortho-cleavage, protocatechuate ortho- and meta-cleavage, and N-acquisition pathways alongside the species tree generated with Orthofinder using the plotTree.barplot function of Phytools (https://cran.r-project.org/web/packages/phytools/index.html) in R v.4.3.2^63^.

To test enrichment of gene families in pyrophilous versus non-pyrophilous isolates, we implemented generalized linear models (GLMs) with a Poisson distribution using lme4 package in R^64^, using gene family counts as the response variable, fungal lifestyle (pyrophilous or non-pyrophilous) as the predictor, and proteome size as an offset. Significance of enrichment was assessed from the trait (pyrophilous vs. non-pyrophilous) coefficient, and false discovery rate (FDR) was controlled using the Benjamini–Hochberg method.

### Additional bioinformatics analysis

We evaluated different mechanisms of gene evolution by examining the ortho groups with aromatic C and N acquisition genes and examining the presence and absence and length of predicted protein amino acid sequences in pyrophilous vs. sister isolates. A gene was labeled core, when it existed in same size in all examined isolates; present/absent when it was either present or absent in the pyrophilous isolate vs. others; fragmented where there were multiple genes in the pyrophilous isolate such that sum of their predicted protein sizes was nearly equal to the whole protein in the sister isolate; and truncated if the protein size was 20% lower than in the sister isolates.

The genome coverage was calculated by QUAST software v.5.2.0^49^ and averaged over 1000 bp windows for visualization by Tinycov v.0.4.0 (https://github.com/cmdoret/tinycov) with the alignment of HiFi reads to the corresponding genome obtained with Minimap2 v.2.30^65^ as an input. Genome-scale alignment of DNA sequences was obtained and visualized by JupiterPlot v.1.1 (https://github.com/JustinChu/JupiterPlot). We measured pair-wise average nucleotide identity between pyrophilous and sister isolates using Jspecies v.1.2.1^66^.

We assessed genome-wide gene fragmentation events by listing tandem genes in *P. simplex* P-BS and P-BF (pyrophilous isolates) that had lower numbers of the corresponding orthologue in *P. simplex* WH6 (non-pyrophilous isolate). For each of loci with fragmented genes, we then calculated the number of genes in three *P. simplex* isolates and visualized it as a circos plot using ShinyCircos v1.0^67^. We also examined if fragmented genes had different transcriptions by visualizing the transcript per million (TPM) value calculated for each gene using TPMcalculator^68^ projected into other layers of the circos plot. We also conducted motif enrichment in the intergenic sequences between fragmented gene derivatives using MEME v.5.5.8^69^. The homologue sequences corresponding to top 5 enriched DNA motifs were retrieved using Tomtom^70^ included with the MEME suit.

We visualized plasmids and their corresponding gene positions using AngularPlasmid v.1.0.5 (https://github.com/vixis/angularplasmid). We identified candidate genomic loci with introgression of aromatic C acquisition and glutamine biosynthesis genes on plasmids by aligning the corresponding plasmid region to the genomes of 16 pyrophilous bacteria using Minimap2 v.2.30^65^ with -cx asm5 parameter. The candidate introgression sites were then visualized using Lovis4u (https://github.com/art-egorov/lovis4u). All multiple protein sequence alignments were conducted using MAFFT v.7^71^ and phylogenetic trees were generated using FastTree v2.1.11^61^ software with -spr 4, -mlacc 2, -slownni, and -gamma parameters. All phylogenetic trees and corresponding data were visualized using iTOL software v.7 (https://itol.embl.de/).

### Acidification and citrate usage by *P. simplex* P-BS and *Pseudomonas* sp

We assessed the media acidifying ability of *P. simplex* P-BS and P-BF and *Pseudomonas* sp. by growing the bacteria on oxidation-fermentation (OF) media (per 1 L distilled water: peptone 2 g, sodium chloride 5 g, dipotassium phosphate 0.3 g, agar 3 g, glucose 10 g [added post autoclave], bromothymol blue 0.03 g)^72^ for 48 h and examining the color change from blue to yellow as indicator of pH reduction. We measured the acidifying strength of these isolates with 6 biological replicates per isolate by growing them in OF broth (all ingredients except for agar) for 48 h and measuring pH with an electrode. Data was analyzed using a linear mixed-effects model (lme4 package in R)^64^, with pH as the response variable, isolate as a fixed effect, and biological replicate as the random effect. Pairwise mean comparisons were performed with Tukey’s HSD adjustment in the emmeans^73^ R package.

We also assessed the isolate’s ability to utilize citrate as a sole carbon source using Simmons’ citrate agar (per 1 L distilled water: sodium citrate 2□g, ammonium dihydrogen phosphate 1□g, dipotassium phosphate 1□g, magnesium sulfate 0.2□g, sodium chloride 0.5□g, ferric ammonium citrate 0.1□g, agar 15□g, and bromothymol blue 0.08□g as a pH indicator)^74^. Isolates were streaked onto the Simmons’ citrate agar and incubated at room temperature for up to 7 days. Growth on the medium with a color change of the indicator from green to blue was recorded as a positive result for citrate utilization.

## Acknowledgements

This project was funded by the Department of Energy BER Award #DE-SC0023127 to SIG, PMH, SDA, and MJW and United States Department of Agriculture-NIFA Award #2022-67014-36675 to SIG and PMH. The RNA-seq data was generated by the U.S. Department of Energy Joint Genome Institute (JGI) under Community Sequencing Project Proposal 510601 to SIG and MJW. The JGI (https://ror.org/04xm1d337) is a DOE Office of Science User Facility supported by the Office of Science of the U.S. Department of Energy and operated under Contract No. DE-AC02-05CH11231. We thank Alex Chase for thoughtful feedback on the manuscript. We thank many individuals who contributed to the Glassman lab pyrophilous bacterial culture collection including former lab managers Judy Chung and James Randolph, lab assistants Ryan Quaal and Aishwarya Veerabahu, UCR Ph.D. students Fabiola Pulido Barriga, Arik Joukhajian, and Mark Yacoub, and UCR undergraduates Anna Nguyen, Marely Vega, Jenna Maddox, Justin Diab, Wine-Jie Lipardo, Audrey Reichard, Priscilla Shultz, Jorge Ponce, and Nathan Heger.

