## SupplementalFiguresandTables for "Genomic innovation through fragmentation and gene transfer underlies adaptation in pyrophilous bacteria"

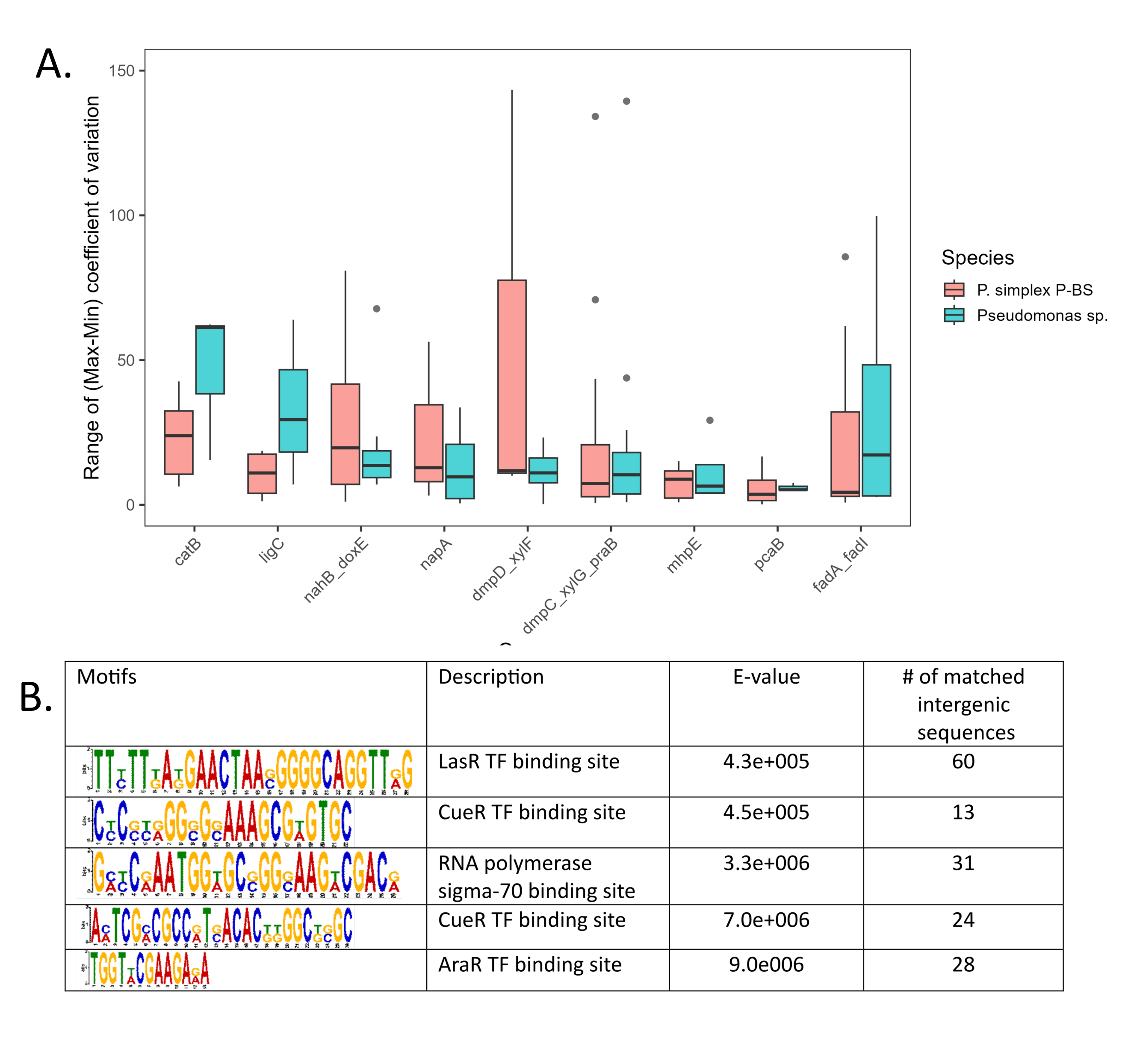

**Fig S1.** Transcriptional variation and intergenic motif enrichment in fragmented aromatic carbon (C) acquisition genes. (A) Variation in transcription of selected aromatic C acquisition genes derived from gene-fragmentation events. Transcriptional variation was calculated as the coefficient of variation (CV) of Transcription Per Million (TPM) values from RNA-seq samples collected at 0, 1, 2, 4, and 16 h post-PyOM treatment for each orthologue. Variation among fragmented derivative orthologues was quantified as the difference between maximum and minimum CV values and summarized per gene family using box plots. Boxes represent the interquartile range (Q1–Q3), the line inside shows the median, whiskers extend to values within 1.5× the interquartile range, and points beyond the whiskers indicate outliers. (B) Motif enrichment in intergenic sequences between fragmented gene derivatives. MEME software was used to identify statistically significant sequence motifs, visualized as sequence logos showing conserved nucleotides and their relative frequencies. Motifs were annotated using Tomtom; E-values indicate the expected number of false positive matches by chance, with lower values denoting more significant similarity. The number of matched intergenic sequences indicates how many sequences contain the identified motif.

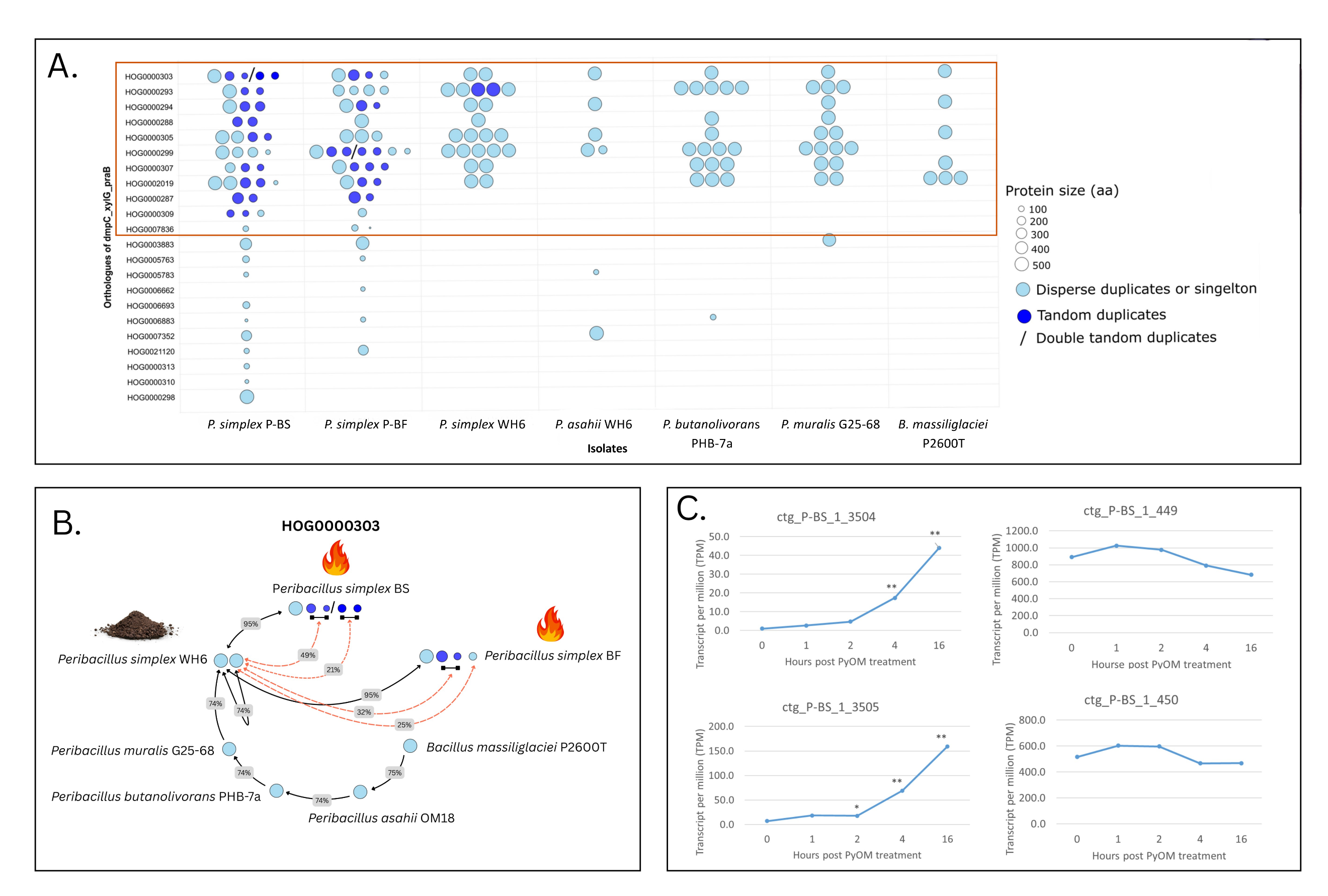

**Fig S2.** The impact of gene fragmentation on the predicted proteome of Aminomuconate-semialdehyde orthologues (*dmpC_xylG_praB*) in pyrophilous isolates *Peribacillus simplex* P-BS and P-BF and their 5 non-pyrophilous sister species (A), the impact of fragmentation on *dmpC_xylG_praB* orthologue HOG0000303 as an example (B) and transcription of fragmented HOG0000303 orthologues in P-BS in response to PyOM treatment (C). The circles in (A) represent the predicted proteome of each gene and their size represents the amino acid length. The brown box shows 11 orthogroups with fragmentation. The black arrows in (B) connect the orthologues in most related isolate among *Peribacillus* species used for orthogroup constructions. The numbers on arrows represent amino acid sequence identity. The dashed red arrows connected the fragmented derivative genes in P-BS and P-BF to their corresponding protein in isolate WH6. Soil and fire icons indicate the origin of isolates being examined. The blackline under circles shows the fragmented gene derivatives of a single gene. Asterisks (P<0.05 *, P <0.01 **) in (C) denote significant differences in transcription compared with 0 h post PyOM exposure, where PyOM significantly induced the transcription of two out of four fragmented gene derivatives.

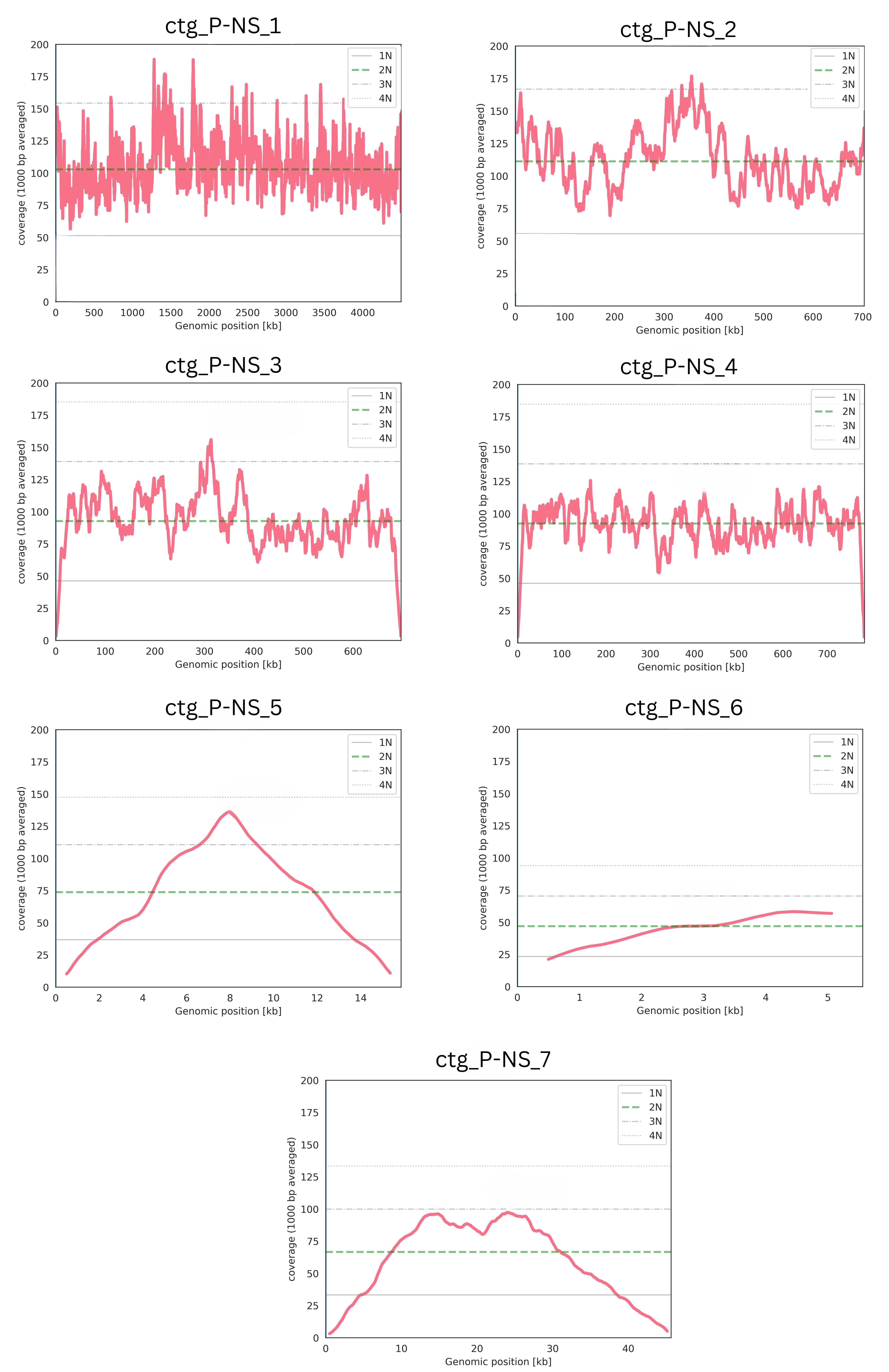

**Fig S3.** Read coverage of *Noviherbaspirillum soli* genome (*ctg_P-NS_1*) compared with 6 candidate plasmids (sizes in Kb: 703, 697, 785, 16, 5, 45). Red lines indicate read coverage averaged over 1,000 bp windows. Horizontal lines represent coverage levels grouped into four quartile ranges (from 1× to 2× upper and lower of the median coverage), and the green dashed line marks the median coverage.

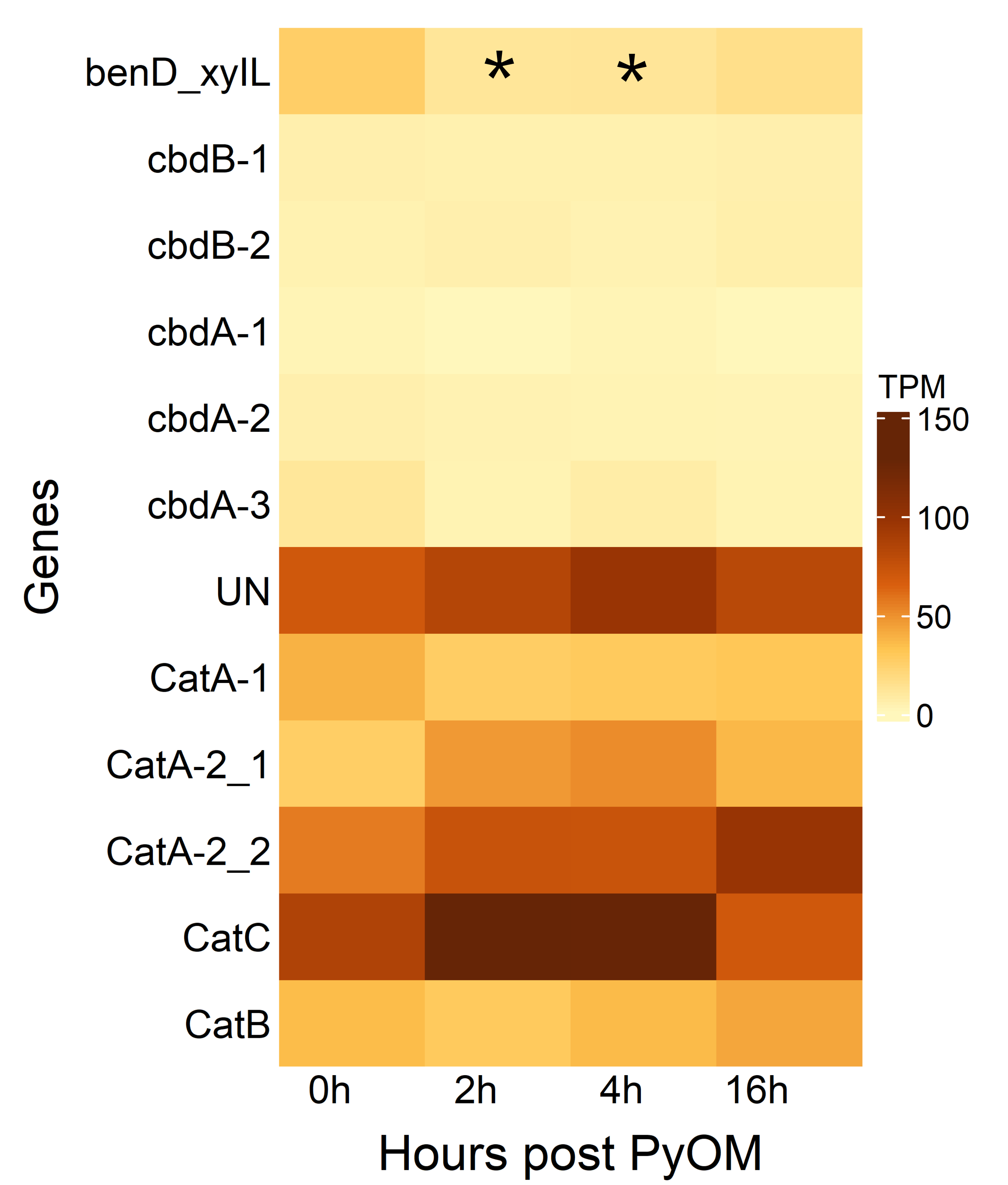

**Fig S4.** Transcription of *Pseudomonas* sp. genes in a region with putative horizontal gene transfer from the *Noviherbaspirillum soli* beta-ketoadipate pathway (BKA) operon on plasmid *ctg P-NS 4*. Gene symbols correspond to those in Fig. 3C. PyOM denotes a medium prepared from charcoal as described in Methods. TPM, transcripts per million. Asterisks (*) indicate significant differences (P< 0.05) relative to 0 h post-PyOM exposure. Transcription analysis revealed that only a gene encoding cis-dihydrobenzoate dehydrogenase (*benD-xyIL*) that had homology with *benc* on the *N. soli* plasmid was significantly induced post PyOM exposure and regions with transposon insertions (*cdb* genes, marked with arrows in Fig. 3C) exhibited lower transcription than adjacent *Cat* genes without insertions.

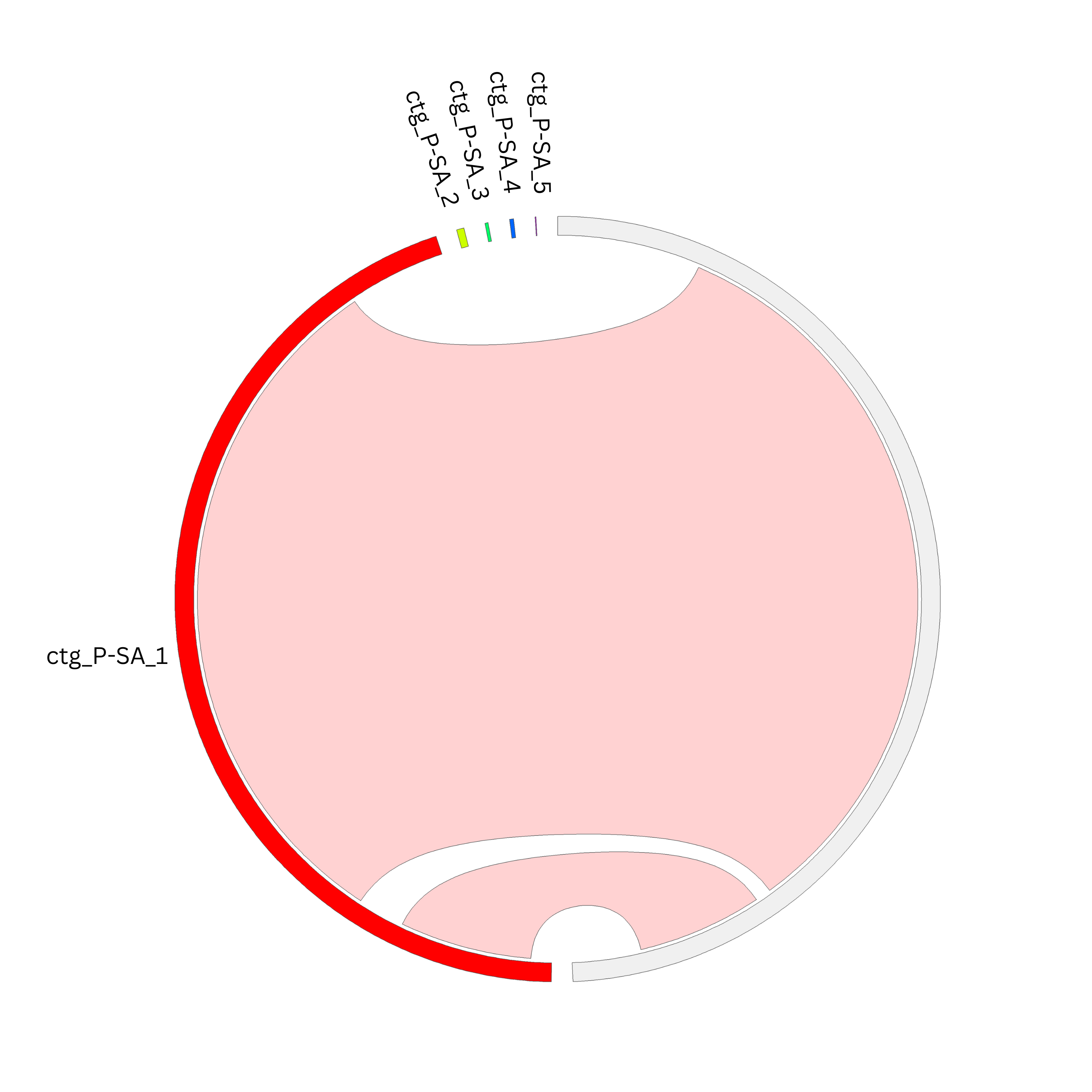

**Fig S5.** Genome and plasmid alignments of the pyrophilous *Streptomyces longhuiensis* (red bar) with its non-pyrophilous sister isolate *S. longhuiensis* BH-MK-02 (gray bar) showing that plasmids *ctg_P-SA_2, ctg_P-SA_3, ctg_P-SA_4*, and *ctg_P-SA_5* have no alignment (red ribbons) to the sister genome and are unique to the pyrophilous isolate.

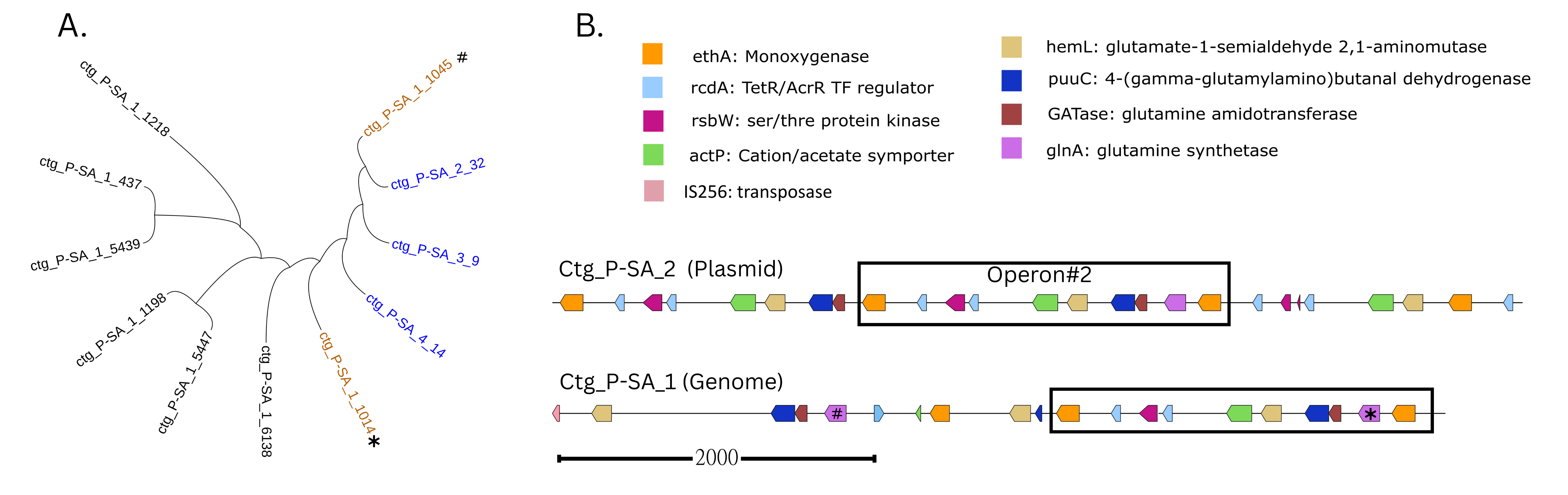

**Fig S6.** Plasmid-mediated operon expansion of glutamine synthetase gene cluster into the *Streptomyces longhuiensis* genome. (A) Phylogeny comparing plasmid (ID in blue) and closely related (ID in brown) and distantly related (ID in black) genomic copies of the operon structural gene, *glnA*. (B) Gene order of candidate expanded genes: operon#2 on plasmid *ctg_P-SA_2* versus the corresponding genomic locus (*ctg_P-SA_1*). The rectangle highlights an intact region transferred from the plasmid to the genome. Scale bar represents 2000 bp. The # and * shows the two genomic *glnA* with homology to plasmid genes in phylogenetic tree in (A) and operon map in (B).

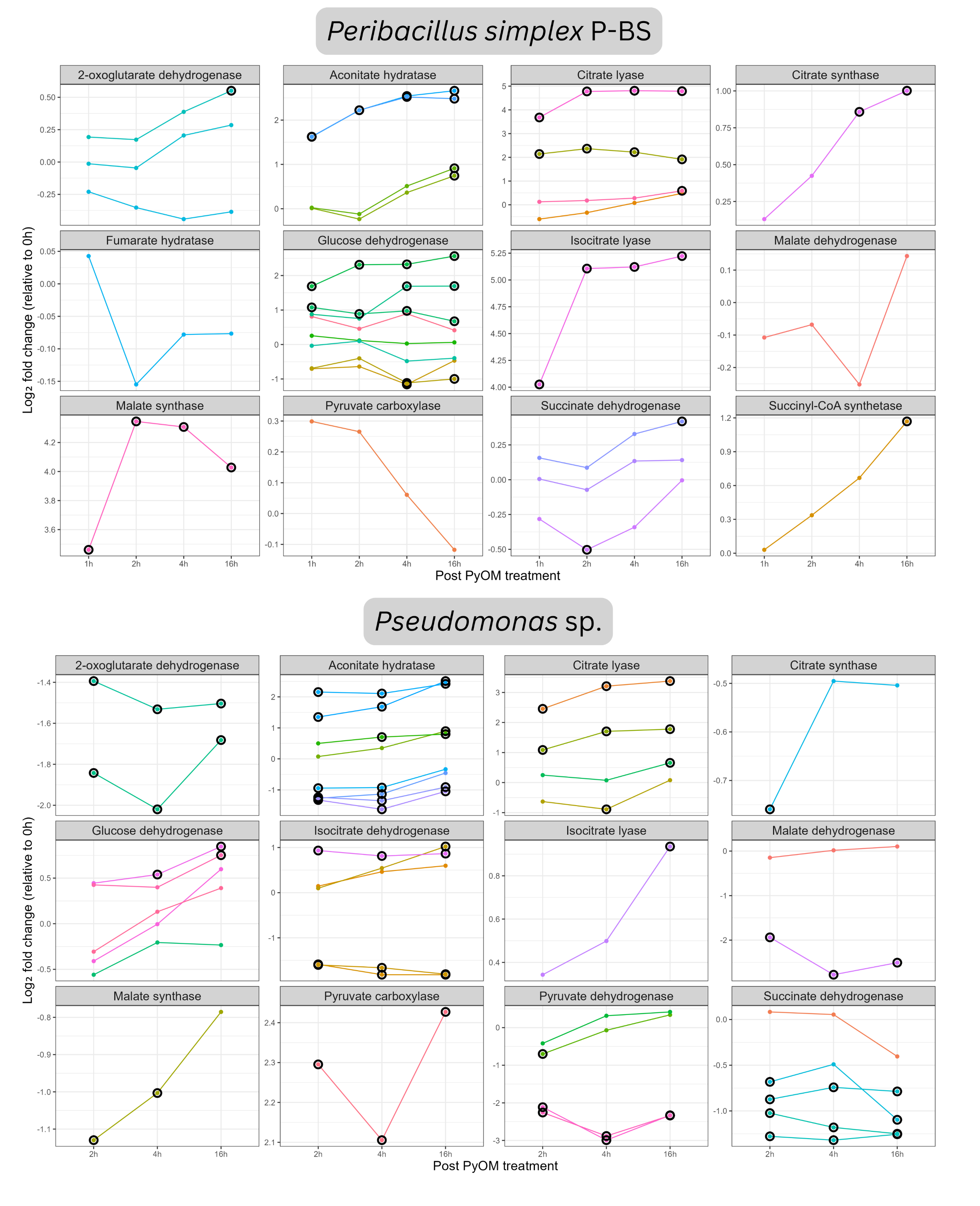

**Fig S7.** Transcriptional response of Tricarboxylic acid (TCA) pathway genes in *Peribacillus simplex* P-BS and *Pseudomonas* sp. Each plot shows single or multiple orthologues of a given TCA pathway gene, represented by individual lines. The log_2_fold change was calculated using DESeq2 from RNA-seq data before, and at 2,4, and 16 h after PyOM (media prepared from charcoal) exposure. Samples collected at 1 h post PyOM for *Pseudomonas* sp. exposure were excluded due to high inter-replicate variations. Black circles indicate significant differences relative to 0 h post-PyOM exposure (P < 0.05).

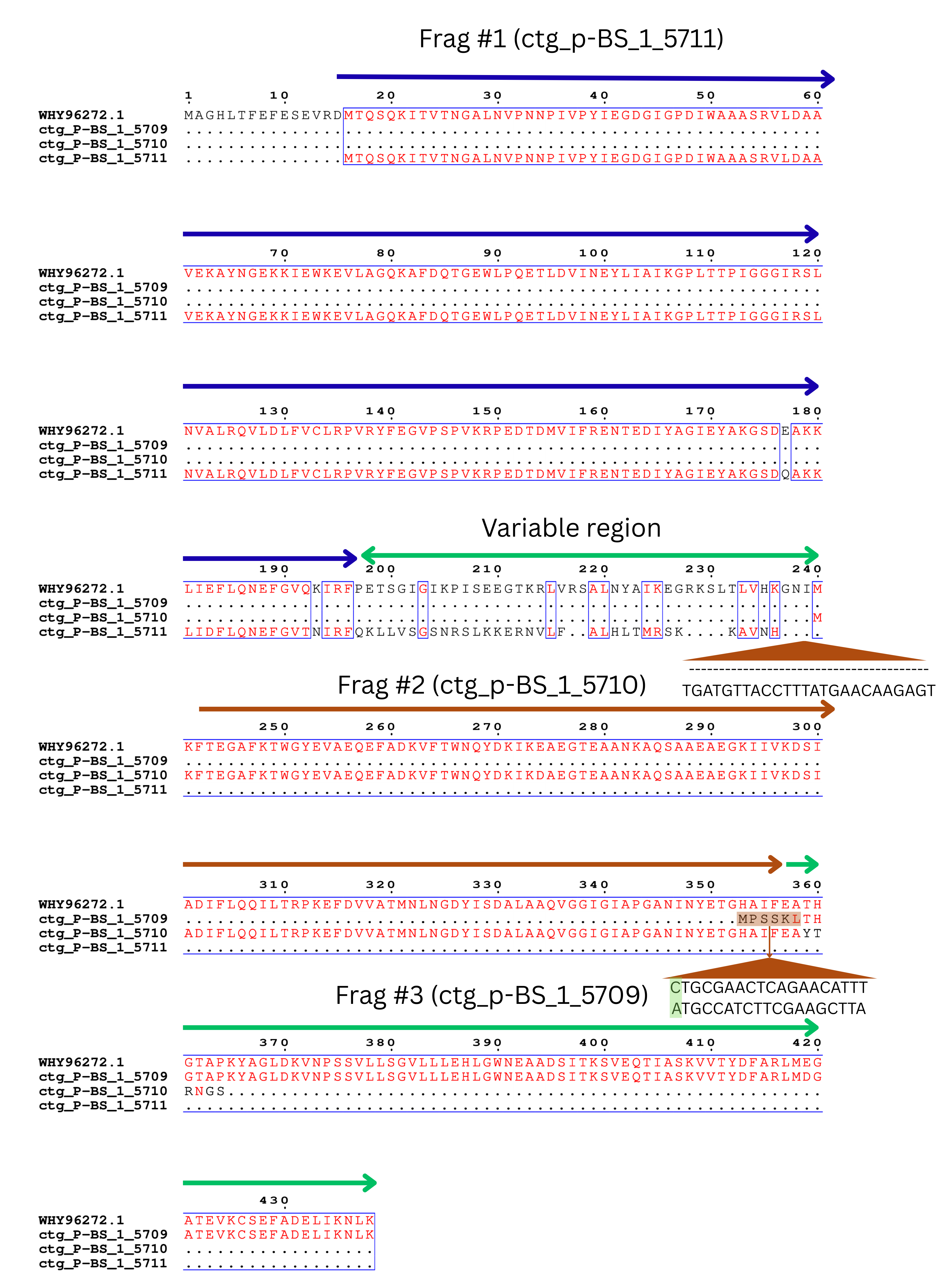

**Fig S8.** Protein sequence alignment of *isocitrate dehydrogenase* (*IDH*) fragmented genes (*ctg_p_BS_1_5709-5711*) from pyrophilous isolate *Peribacillus simplex* P-BS and its homolog in non pyrophilous sister isolate *P. simplex* WH6 (WHY96272.1). Arrows in blue, brown, and green indicate the positions of fragmented *IDH* in P-BS projected onto the WH6 reference. The green region between the brown and blue arrows marks a hypervariable segment separating the two fragments. Brown triangles denote sequence divergences associated with fragmentation, with corresponding nucleotide sequences shown below: the upper triangle indicates a 25-nucleotide insertion between fragments 1 and 2, and the lower triangle (green shading) marks a single-nucleotide polymorphism introducing a novel ATG start codon with the corresponding protein sequence highlighted in the alignment.

**Table S1.** Species names, isolate ID, phylogenetic classifications, wildfire origins, sequencing coverage (calculated by dividing the total length of genome to total number of bases generated through sequencing), assembly metrics including genome size, plasmid number and sizes of those plasmids, and BUSCO coverage, and NCBI project IDs for the 16 isolates of pyrophilous bacteria (8 Actinomycetota, 4 Bacillota, 4 Pseudomonadota). We used PacBio high fidelity (HiFi) sequencing to assemble all bacterial genomes into a single contig. All bacteria were isolated from burned soil or smoke (identified with “‡”) from within or nearby US National Forests in Southern California, USA. SBNF = San Bernadino National Forest; CNF = Cleveland National Forest; ANF = Angeles National Forest. SBNF is within San Bernadino county, CNF is within Riverside and Orange Counties, and ANF is within Los Angeles county. All isolates are stored in glycerol stocks in a -80 °C ultracold freezer in the Glassman lab and were given unique isolate IDs. The BUSCO column reports the percentage of completion (C), with the percentage of duplicated genes indicated in brackets, as well as the percentages of fragmented (F) and missing (M) BUSCO genes of genomes assembled for each isolate.

| **Species** | **Isolate ID** | **Phylum: Class: Order: Family** | **Wildfire name, year, national forest** | **HiFi coverage (X)** | **Genome size (Gb)** | **Number of plasmids and sizes** | **BUSCO** | **NCBI project #** |
| --- | --- | --- | --- | --- | --- | --- | --- | --- |
| *Massilia litorea* | R2A_B_28 | Pseudomonadota: Betaproteobacteria: Burkholderiales: Oxalobacteraceae | El Dorado Fire, 2020, SBNF | 172 | 5.1 | 0 | C:99.3[0.7]; F:0.2; M:0.5 | PRJNA1348574 |
| *Methylobacterium gregans*‡ | B_121 | Pseudomonadota: Alphaproteobacteria: Hyphomicrobiales: Methylobacteriaceae | Holy Fire, 2018, CNF | 133 | 6.0 | 0 | C:99.8 [1.2]; F:0; M:0.2 | PRJNA1348575 |
| *Noviherbaspirillum soli* | PyOM_B_09 | Pseudomonadota:  Betaproteobacteria: Burkholderiales: Oxalobacteraceae | El Dorado Fire, 2020, SBNF | 104 | 4.5 | 6 (703,697,785,16,5,45) | C:100 [0.9]; F:0; M:0 | PRJNA1348585 |
| *Pseudomonas sp.* | 4C_B_06 | Pseudomonadota:Gammaproteobacteria: Pseudomonadales: Pseudomonadaceae | El Dorado Fire, 2020, SBNF | 54 | 6.3 | 0 | C:99.7[0]; F:0; M:0.3 | PRJNA1348587 |
| *Arthrobacter globiformis* | T4 B01 | Actinomycetota: Actinomycetia: Micrococcales: Micrococcaceae | El Dorado Fire, 2020, SBNF | 102 | 5.1 | 3 (301,44,7) | C:99.3[1.4]; F:0; M:0.7 | PRJNA1348183 |
| *Arthrobacter ipis* | T4 B16 | Actinomycetota: Actinomycetia: Micrococcales: Micrococcaceae | El Dorado Fire, 2020, SBNF | 100 | 4.8 | 1 (7) | C:99.3[0.7]; F:0; M:0.7 | PRJNA1348207 |
| *Aermicrobium* sp. P-AF | 4C_B_16 | Actinomycetota: Actinomycetia: Propionibacteriales: Nocardioidaceae | El Dorado Fire, 2020, SBNF | 103 | 3.4 | 0 | C:99.7[0]; F:0; M:0.3 | PRJNA1348175 |
| *Aermicrobium* sp. P-AG | R2A_B_31 | Actinomycetota: Actinomycetia: Propionibacteriales: Nocardioidaceae | El Dorado Fire, 2020, SBNF | 177 | 3.3 | 0 | C:99.7[0]; F:0; M:0.3 | PRJNA1348180 |
| *Kocuria salina* | FRZ_B_22 | Actinomycetota: Actinomycetia: Micrococcales: Micrococcaceae | El Dorado Fire, 2020, SBNF | 60 | 3.8 | 2 (107,6) | C:98.9[0.3]; F:0.7; M:0.4 | PRJNA1348526 |
| *Micrococus yunnanensis*‡ | BS_B04 | Actinomycetota: Actinomycetia: Micrococcales: Micrococcaceae | Bobcat Fire, 2020, ANF | 77 | 2.5 | 0 | C:98.3[0]; F:0.3; M:1.4 | PRJNA1348577 |
| *Modestobacter versicolor‡* | B_119 | Actinomycetota: Actinomycetia: Geodermatophilales: Modestobacteraceae | Holy Fire, 2018, CNF | 142 | 4.1 | 1 (663) | C:99.6[1]; F:0; M:0.4 | PRJNA1348579 |
| *Streptomyces* *longhuiensis* | DRY_B_16 | Actinomycetota: Actinomycetia: Streptomycetales: Streptomycetaceae: | El Dorado Fire, 2020, SBNF | 37 | 8.9 | 4 (62,26,34,7) | C:99.9[0.8]; F:0.1; M:0 | PRJNA1348588 |
| *Bacillus pumillus*‡ | BS_B01 | Bacillota: Bacilli; Bacillales: Bacillaceae | Bobcat Fire, 2020, ANF | 358 | 3.7 | 1 (10) | C:100[1.3]; F:0; M:0 | PRJNA1348485 |
| *Peribaciilus simplex* P-BF | PyOM_B_05 | Bacillota: Bacilli; Bacillales: Bacillaceae | El Dorado Fire, 2020, SBNF, San Bernardino | 241 | 5.8 | 0 | C:99.7[0]; F:0; M:0.3 | PRJNA1348474 |
| *Peribaciilus simplex* P-BS | 4C_B_15 | Bacillota: Bacilli; Bacillales: Bacillaceae | El Dorado Fire, 2020, SBNF | 254 | 5.6 | 0 | C:100[0]; F:0; M:0 | PRJNA1348524 |
| *Priestia aryabhattai* | B_39 | Bacillota: Bacilli; Bacillales: Bacillaceae | Holy Fire, 2018, CNF | 273 | 5.2 | 2 (10,7) | C:99.7[1]; F:0.3; M:0 | PRJNA1348476 |

**Table S2.** The public genomes used for orthogroup construction, their gene bank accessions and their contig N50.

| **Isolates** | **GenBank accession** | **contigs N50 (Kb)** |
| --- | --- | --- |
| *Aeromicrobium fastidiosum* DSM_10552 | GCF_017876595.1 | 306 |
| *Aeromicrobium ginsengisoli* JCM_14732 | GCF_004134905.2 | 122 |
| *Aeromicrobium marinum* DSM_15272 | GCF_000160775.2 | 191 |
| *Aeromicrobium massiliense* JC14 | GCF_000312105.1 | 4,826 |
| *Arthrobacter globiformis* V3I9 | GCF_030817195.1 | 5,568 |
| *Arthrobacter ipis* IA7 | GCF_014694315.1 | 4,919 |
| *Bacillus australimaris* NH7I_1 | GCF_001307105.1 | 5,786 |
| *Bacillus massiliglaciei* P2600T | GCF_900098925.1 | 5,267 |
| *Bacillus pumilus* 145 | GCA_003431975.1 | 4,000 |
| *Bacillus safensis* U14-5 | GCA_001938665.1 | 4,559 |
| *Bacillus zhangzhouensis* SC123 | GCF_031195075.1 | 4,963 |
| *Kocuria rosea* ATCC_186 | GCA_006094695.1 | 5,044 |
| *Kocuria rosea* CMS_76or | GCF_000786195.1 | 5,986 |
| *Kocuria salina* CV6 | GCF_013368965.1 | 5,521 |
| *Kocuria sediminis* JCM_17929 | GCF_009735315.1 | 2,839 |
| *Massilia brevitalea* CGMCC_1.10731 | GCF_030711405.1 | 6,551 |
| *Massilia litorea* LPB0304 | GCA_015101885.1 | 9,817 |
| *Massilia yuzhufengensis* CGMCC_1.12041 | GCF_900112225.1 | 6,849 |
| *Methylobacterium dankookense* SW08-7 | GCF_902141855.1 | 8,615 |
| *Methylobacterium gregans* DSM_19564 | GCF_030814915.1 | 7,868 |
| *Methylobacterium hispanicum* DSM_16372 | GCF_022179285.1 | 306 |
| *Methylobacterium radiodurans* 17Sr1-43 | GCA_003173735.1 | 122 |
| *Micrococcus aloeverae* RAT4 | GCF_014138395.1 | 191 |
| *Micrococcus luteus* ML | GCA_024397455.1 | 4,826 |
| *Micrococcus yunnanensis* TT9 | GCA_023573625.1 | 5,568 |
| *Modestobacter altitudinis* 1G4 | GCF_005930475.1 | 4,919 |
| *Modestobacter italicus* BC_501 | GCA_000306785.1 | 5,786 |
| *Modestobacter versicolor* DSM_16678 | GCF_014195485.1 | 5,267 |
| *Noviherbaspirillum pedocola* DKR-6 | GCF_016632335.1 | 4,000 |
| *Noviherbaspirillum soli* SUEMI10 | GCF_015352955.1 | 4,559 |
| *Noviherbaspirillum suwonense* DSM_26001 | GCF_900182955.1 | 4,963 |
| *Peribacillus asahii* OM18 | GCA_004006295.1 | 5,044 |
| *Peribacillus butanolivorans* PHB-7a | GCA_003410415.1 | 5,986 |
| *Peribacillus muralis* G25-68 | GCA_001645685.2 | 5,521 |
| *Peribacillus simplex* WH6 | GCA_030123485.1 | 2,839 |
| *Priestia aryabhattai* KNUC0119 | GCF_018881345.1 | 6,551 |
| *Priestia flexa* DMP08 | GCA_021441905.1 | 9,817 |
| *Priestia koreensis* FS-1 | GCA_022646885.1 | 6,849 |
| *Pseudarthrobacter psychrotolerans* YJ56 | GCA_009911795.1 | 8,615 |
| *Pseudarthrobacter sulfonivorans* Ar51 | GCA_001484605.1 | 7,868 |
| *Pseudomonas cichorii* JBC1 | GCA_000517305.1 | 306 |
| *Pseudomonas graminis* PgKB30 | GCA_013201545.1 | 122 |
| *Pseudomonas lutea* DSM_17257 | GCF_000759445.1 | 191 |
| *Pseudomonas phytophila* ICMP_23753 | GCA_025643095.1 | 4,826 |
| *Streptomyces longhuiensis* BH-MK-02 | GCA_020616555.1 | 5,568 |
| *Streptomyces rutgersensis* NBH77 | GCA_014216335.1 | 9,817 |
| *Streptomyces tsukubensis* AT3 | GCA_009296025.1 | 6,849 |
| *Streptomyces xanthii* CRXT-Y-14 | GCA_014621695.1 | 8,615 |

**Table S3.** Genes involved in multiple pathways of aromatic carbon (C)† degradation from PyOM, and those in acquiring inorganic nitrogen (N) generated during combustion, and tricarboxylic acid (TCA) cycle, and Kyoto Encyclopedia of Genes and Genomes (KEGG) IDs used for annotation of the genes in pyrophilous bacteria and their sister species. For TCA cycle genes, the description includes the reaction in parentheses. There are multiple KEGG IDs when there are multiple isoforms of the enzyme, denoted in parentheses.

| **Gene** | **Description** | **Pathways** | **#KEGG IDs** |
| --- | --- | --- | --- |
| dmpC_xylG_praB | Aminomuconate-semialdehyde/2-hydroxymuconate-6-semialdehyde dehydrogenase | Benzoate degradation† | K10217 |
| *ligC* | 2-hydroxy-4-carboxymuconate | Benzoate degradation† | K10219 |
| *ligI* | 2-pyrone-4,6-dicarboxylate lactonase | Benzoate degradation† | K10221 |
| *pcaI* | 3-oxoadipate CoA-transferase | Benzoate degradation† | K01031 |
| *ligJ* | 4-oxalmesaconate hydratase | Benzoate degradation† | K10220 |
| *galB* | 4-oxalomesaconate hydratase | Benzoate degradation† | K16515 |
| *galD* | 4-oxalomesaconate tautomeras | Benzoate degradation† | K16514 |
| *pcaL_pcaD* | 3-oxoadipate-enol-lactonase | BKA Pathway† | K01055 |
| *fadA_fadI* | Acetyl-Co A acyltransferase | BKA Pathway† | K00632 |
| *dmpD_xylF* | 2-hydroxymuconate-semialdehyde hydrolase | Catechol Metacleavage† | K10216 |
| *dmpH_xylI* | 2-oxo-3-hexenedioate decarboxylase | Catechol Metacleavage† | K01617 |
| *praC_xylH* | 4-oxalocrotonate_tautomerase | Catechol Metacleavage† | K01821 |
| *mhpF* | Acetaldehyde_dehydrogenase | Catechol Metacleavage† | K04073 |
| *dmpB_xylE* | Catechol 2,3-dioxygenase | Catechol Metacleavage† | K00446 |
| *mhpE* | Hydroxy_oxovalerate_aldolase | Catechol Metacleavage† | K01666 |
| *pcaJ* | 3-oxoadipate enol-lactonase | Catechol Orthocleavage† | K01032 |
| *catA* | Catechol-1,2-dioxygenase | Catechol Orthocleavage† | K03381 |
| *catB* | Muconate cycloisomerase | Catechol Orthocleavage† | K01856 |
| *catC* | Muconolactone D-isomerase | Catechol Orthocleavage† | K03464 |
| *nahD* | 2-hydroxychromene-2-carboxylate-isomerase | Naphthalene degredation† | K14584 |
| *nahB_doxE* | Cis-1,2-dihydro-1,2-dihydroxy naphthalene | Naphthalene degredation† | K14582 |
| *nahA* | Naphthalene-1,2-dioxygenase subunit-alpha | Naphthalene degredation† | K14579 |
| *nahE* | Trans-o-hydroxybenzylidenepyruvate | Naphthalene degredation† | K14585 |
| *ligK_galC* | 4-hydroxy-4-methyl-2-oxoglutarate aldolase | PCA Metacleavage† | K10218 |
| *ligA_LigB* | Protocatechuate-dioxygenase | PCA Metacleavage† | K04100 |
| *pcaB* | 3-carboxy-cis,cis-muconate-cycloisomerase | PCA Orthocleavage† | K01857 |
| *pcaH_pcaG* | Protocatechuate-3,4-dioxygenase | PCA Orthocleavage | K00448 |
| *napA* | Nitrate reductase | N acquisition | K02567 |
| *narG_narZ* | Nitrate-reductase | N acquisition | K00370 |
| *nirK* | Nitrite-reductase | N acquisition | K00368 |
| *nifH* | Nitrogenase iron protein NifH | N acquisition | K02588 |
| *CitC* | Citrate synthase (Oxaloacetate + Acetyl-CoA→Citrate) | TCA cycle | K01647 |
| *CitE* | Citrate lyase (Citrate→Acetyl-CoA) | TCA cycle | K01643 (alpha), K01644 (beta), K01646 (gamma) |
| *ACO* | Aconitate hydratase (Citrate→Isocitrate) | TCA cycle | K01681 |
| *IDH* | Isocitrate dehydrogenase (Isocitrate→α-ketoglutarate) | TCA cycle | K00030 |
| *OGDH* | 2-oxoglutarate dehydrogenase (α-ketoglutarate→Succinyl-CoA | TCA cycle | K00164 (E1), K00658 (E2), K00382 (E3) |
| *suc* | Succinyl-CoA synthetase (Succinyl-CoA ⇌Succinate) | TCA cycle | K01899 (alpha), K01900 (beta-ADP), K01901 (beta-GDP) |
| *SDH* | Succinate dehydrogenase (Succinate→Fumarate) | TCA cycle | K00239 (flavoprotein), K00240 (iron-sulfur) |
| *frd* | Fumarate hydratase (Fumarate ⇌L-malate) | TCA cycle | K01679 |
| *MDH* | Malate dehydrogenase (L-malate→Oxaloacetate) | TCA cycle | K00025 |
| *PC* | pyruvate carboxylase (Pyruvate →Oxaloacetate) | TCA cycle | K01958 |
| *PDHA* | pyruvate dehydrogenase (Pyruvate→Acetyl-CoA) | TCA cycle | K00161 |
| *aceA* | Isocitrate lyase (Isocitrate→Succinate+Glyoxylate) | TCA cycle-glyoxylate shunt | K01637 |
| *aceB* | Malate synthase (Glyoxylate + Acetyl-CoA →Malate + CoA-SH) | TCA cycle-glyoxylate shunt | K01638 |
